# Microbial association networks in cheese: a meta-analysis

**DOI:** 10.1101/2021.07.21.453196

**Authors:** Eugenio Parente, Teresa Zotta, Annamaria Ricciardi

## Abstract

Interactions among starter and non-starter microorganisms (starter bacteria, naturally occurring or intentionally added non-starter bacteria, yeasts and filamentous fungi, spoilage and pathogenic microorganisms and, finally bacteriophages and even arthropods) deeply affect the dynamics of cheese microbial communities and, as a consequence, multiple aspects of cheese quality, from metabolites affecting the taste, aroma and flavour, to body, texture and colour. Understanding and exploiting microbial interactions is therefore key to managing cheese quality. This is true for the simplest systems (fresh cheeses produced from pasteurized milk using defined starters composed solely of Lactic Acid Bacteria) and the more so for complex, dynamic systems, like surface ripened cheese produced from raw milk, in which a dynamic succession of diverse microorganisms is essential for obtained the desired combination of sensory properties while guaranteeing safety. Positive (commensalism, protocooperation) and negative (competition, amensalism, predation and parasitism) among members of the cheese biota have been reviewed multiple times. Although the complex, multidimensional datasets generated by multi-omic approaches to cheese microbiology and biochemistry are ideally suited for the representation of biotic and metabolic interactions as networks, network science concepts and approaches are rarely applied to cheese microbiology.

In this review we first illustrate concepts relevant to the description of microbial interaction networks using network science concepts. Then, we briefly review methods used for the inference and analysis of microbial association networks and their potential use in the interpretation of the cheese interactome. Since these methods can only be used for mining microbial associations, a review of the experimental methods used to confirm the nature of microbial interactions among cheese microbes. Finally, we demonstrate the potential of microbial association network inference by mining metataxonomic data stored in the public database DairyFMBN, a specialized version of FoodMicrobionet which collates data on 74 metataxonomic studies on dairy products. Microbial association networks were inferred from 34 studies on cheese with up to 4 different methods and the results discussed to evaluate several aspects (choice of method, level of taxonomic resolution for the analysis, network, node and edge properties) which provide insight on the usefulness of this approach as explorative tool in the detection of microbial interactions in cheese.

**Highlights:**

1. Approaches for inference of association networks from metataxonomic data were reviewed
2. A metastudy on association networks in cheese was carried out using 34 recent studies
3. Inference method and taxonomic resolution should be chosen carefully
4. SPIEC-EASI may be used as a conservative method for microbial association inference
5. Edge and node properties support the formulation of testable hypotheses for microbial interactions

## 1. Foreword

Cheeses, like all fermented foods, are man-made dynamic ecosystems, in which the environment is organic and the biota (with a few exceptions in which arthropods play a more or less beneficial role (Carvalho et al., 2020; Marcellino and Benson, 2014) is made solely by microbes (bacteria, fungi and viruses) (Gobbetti et al., 2018; Jonnala et al., 2018; Wolfe and Dutton, 2015). Microbial metabolism is among the main drivers of cheese sensory properties, and the dynamics of the microbiota strongly impacts cheese quality and safety (Gobbetti et al., 2018; Jonnala et al., 2018). Even when the complex microbiota of raw milk is drastically simplified by heat treatments, by the addition of defined starter cultures, and when ripening and storage are relatively short (like in fresh cheeses), microbial interactions (from parasitism, when lytic bacteriophages infect starter strains, to proto-cooperation among key starter species, like *Streptococcus thermophilus* and *Lactobacillus delbrueckii* subsp. *bulgaricus*) (Blaya et al., 2017; Irlinger and Mounier, 2009) are still important in determining the success of the fermentation. On the other side of the spectrum, in raw milk cheeses produced with no starter or by using traditional undefined starters, and with longer ripening, a complex pattern of microbial interactions and a succession of species and strains invariably develops and its control is key to cheese quality (Blaya et al., 2017; Gobbetti et al., 2018; Irlinger and Mounier, 2009; Jonnala et al., 2018; Mayo et al., 2021). Microbial successions and interactions are especially complex in surface ripened cheeses (Irlinger and Mounier, 2009), and this has been demonstrated in an elegant and comprehensive way in a series of recent studies (Bonham et al., 2017; Cosetta et al., 2020; Kastman et al., 2016; Niccum et al., 2020; Wolfe et al., 2014; Zhang et al., 2018).

In all ecosystems, several types of positive (commensalism, proto-cooperation) and negative (competition, amensalism, parasitism) interactions are possible between couple of partners or among more complex modules and cliques (see Canon et al., 2020; D’Souza et al., 2018 for recent reviews). Even if, ultimately, co-culturing in the laboratory and/or in appropriate model systems is the only way to obtain in-depth knowledge on the nature of interactions (Cosetta and Wolfe, 2019; D’Souza et al., 2018; Wolfe et al., 2014), the wealth of data provided by metataxonomic, metagenomic and metabolomics approaches provides ample opportunity to mine for microbial association networks and metabolic networks (Layeghifard et al., 2017; Liu et al., 2020; Röttjers and Faust, 2018).

Microbial interactions in cheese and in other fermented foods have been the subject of recent comprehensive reviews (Blaya et al., 2017; Canon et al., 2020; Gobbetti et al., 2018; Mayo et al., 2021). A schematic, and possibly oversimplified representation of interactions (parasitism, commensalism, amensalism, competition, protocooperation) occurring in an idealized surface ripened cheese is shown in Figure 1, with some of the interactions described in detail in section 1.3 and Table 1.

**Figure 1.**
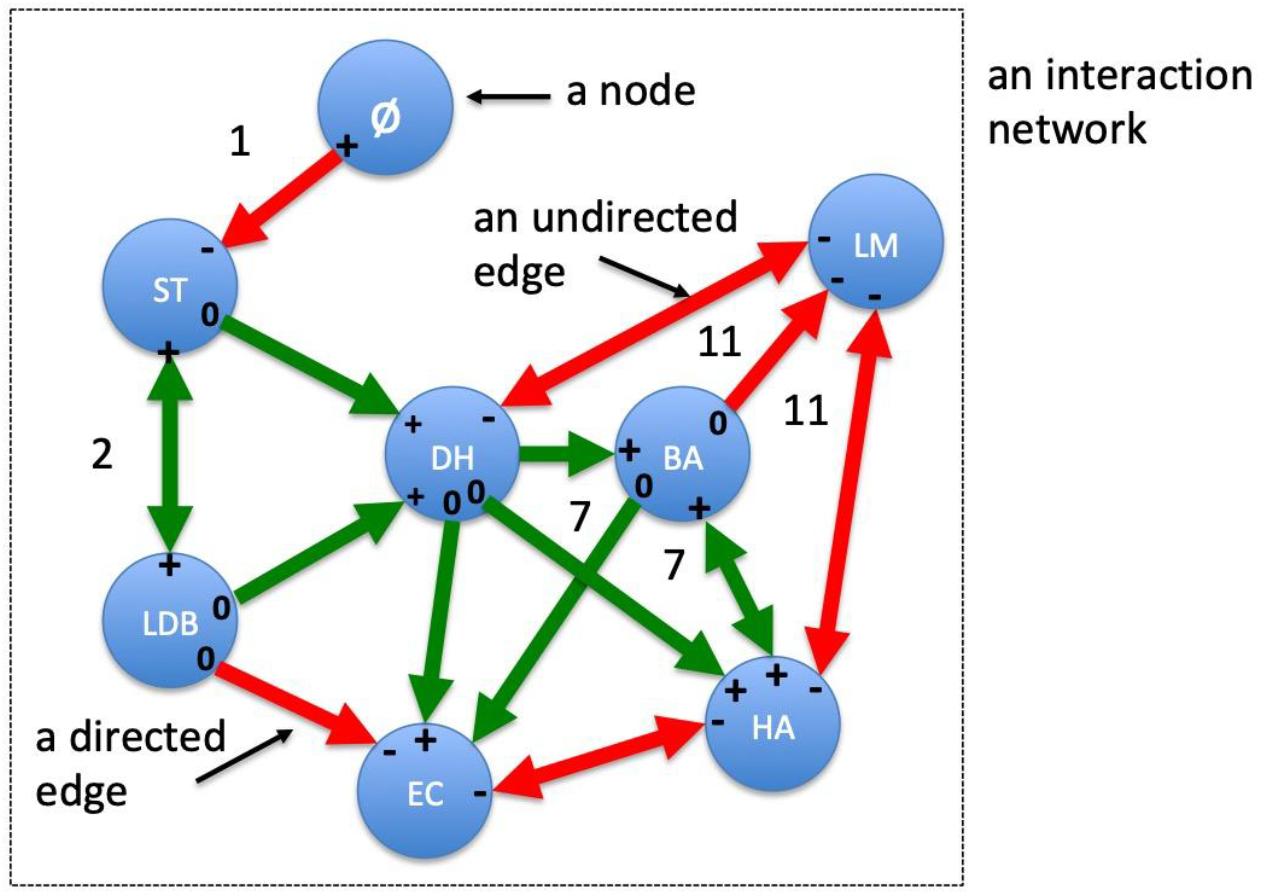
A simplified representation of a potential interaction network in a surface ripened cheese. Red arrows indicate interactions which have a negative effect (often referred to as mutual exclusion interactions) on one or both partners. Green arrows indicate interactions which have a positive effect on one or both partners (often referred to as mutual exclusion interactions). 0 indicates no effect, + a positive effect, - a negative effect. Numbers refer to examples in Table 1. The direction of the arrows may be used to indicate the direction of the interaction. **Parasitism** (1) due to bacteriophage (⌀) infections of starter bacteria, like *Streptococcus thermophilus* (ST) is one of the most frequent, and technologically relevant interactions in cheesemaking, together with the **protocooperation** between starter species (2), like *S. thermophilus* and *Lactobacillus delbrueckii* subsp. *bulgaricus* (LDB). Starter lactic acid bacteria (SLAB) frequently inhibit pathogenic or spoilage bacteria, like *Escherichia coli* (EC) by **competition, amensalism** or **ecosystem conditioning** (decrease of pH, which in turn, due to syneresis and loss of water during ripening, results in reduced a_W_). The latter (decrease in pH due to production of lactic acid, increase in pH due to consumption of lactic acid or proteolysis) is a frequent, indirect type of interaction. Some **commensalistic** (7) relationships (products of the metabolism of one microorganism may become substrate for another) like the use of galactonate produced by the yeast *Debaryomyces hansenii* (DH, which is also responsible of increasing the pH) by *Brevibacterium aurantiacum (BA)* and *Hafnia alvei* (HA). In turn, *H. alvei* may develop commensalistic or proto-cooperative interactions, since siderophores produced by HA stimulate BA, which in turn releases energy compounds for HA from proteins and lipids. Complex cheese consortia are also implicated in the inhibition of *Listeria monocytogenes* (LM) by **amensalism** or **competition** (11).

**Table 1.**
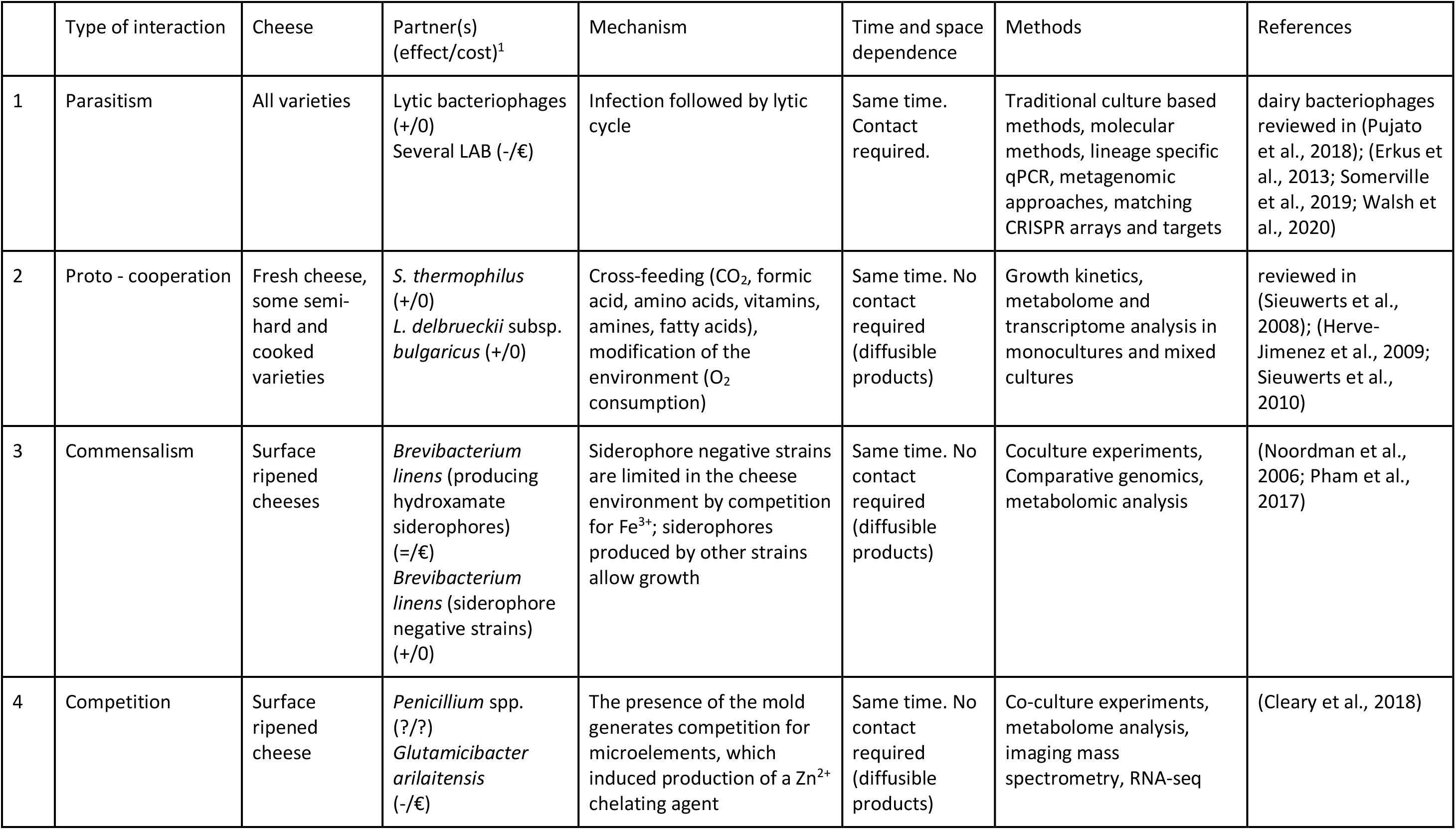
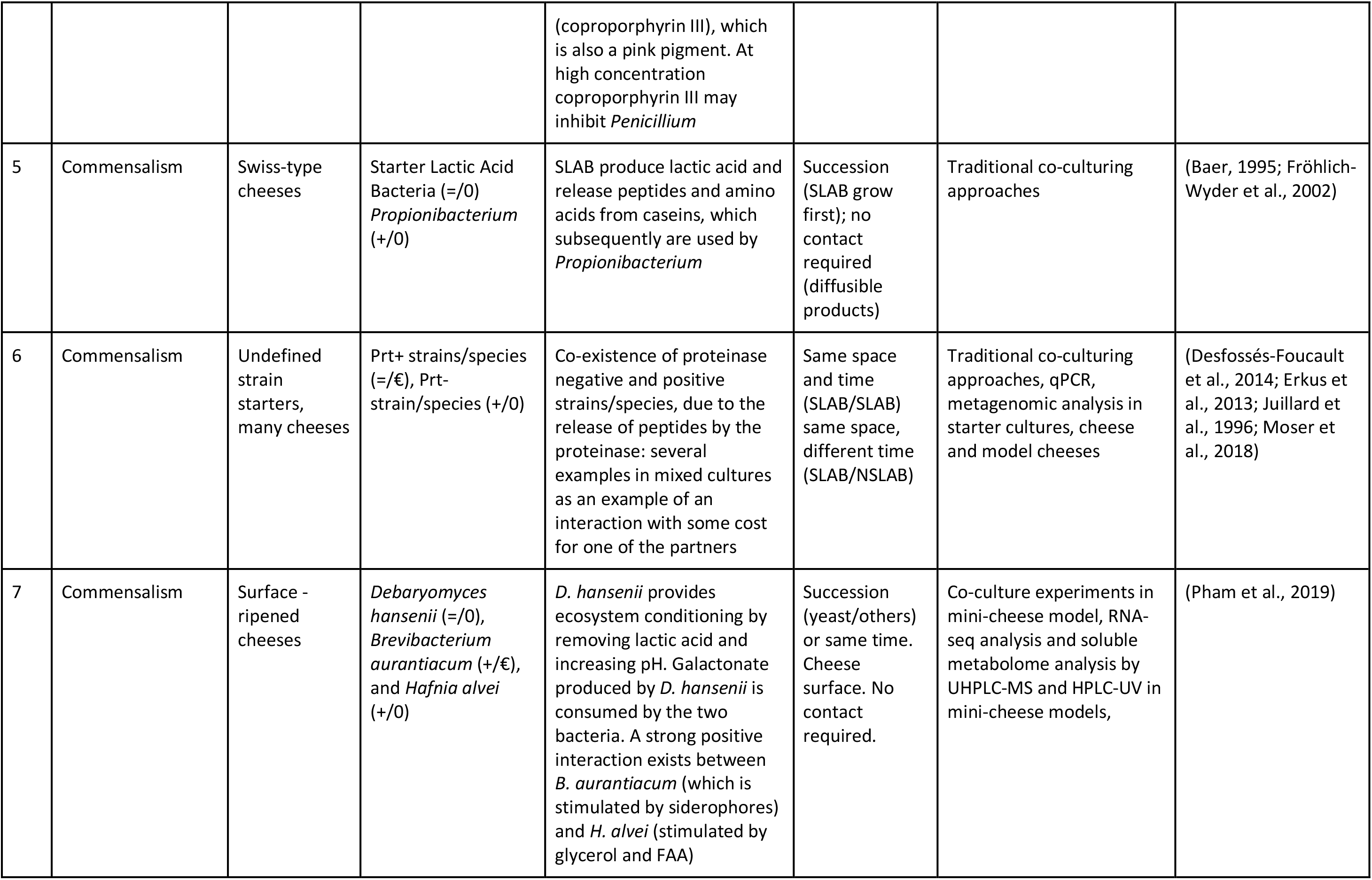
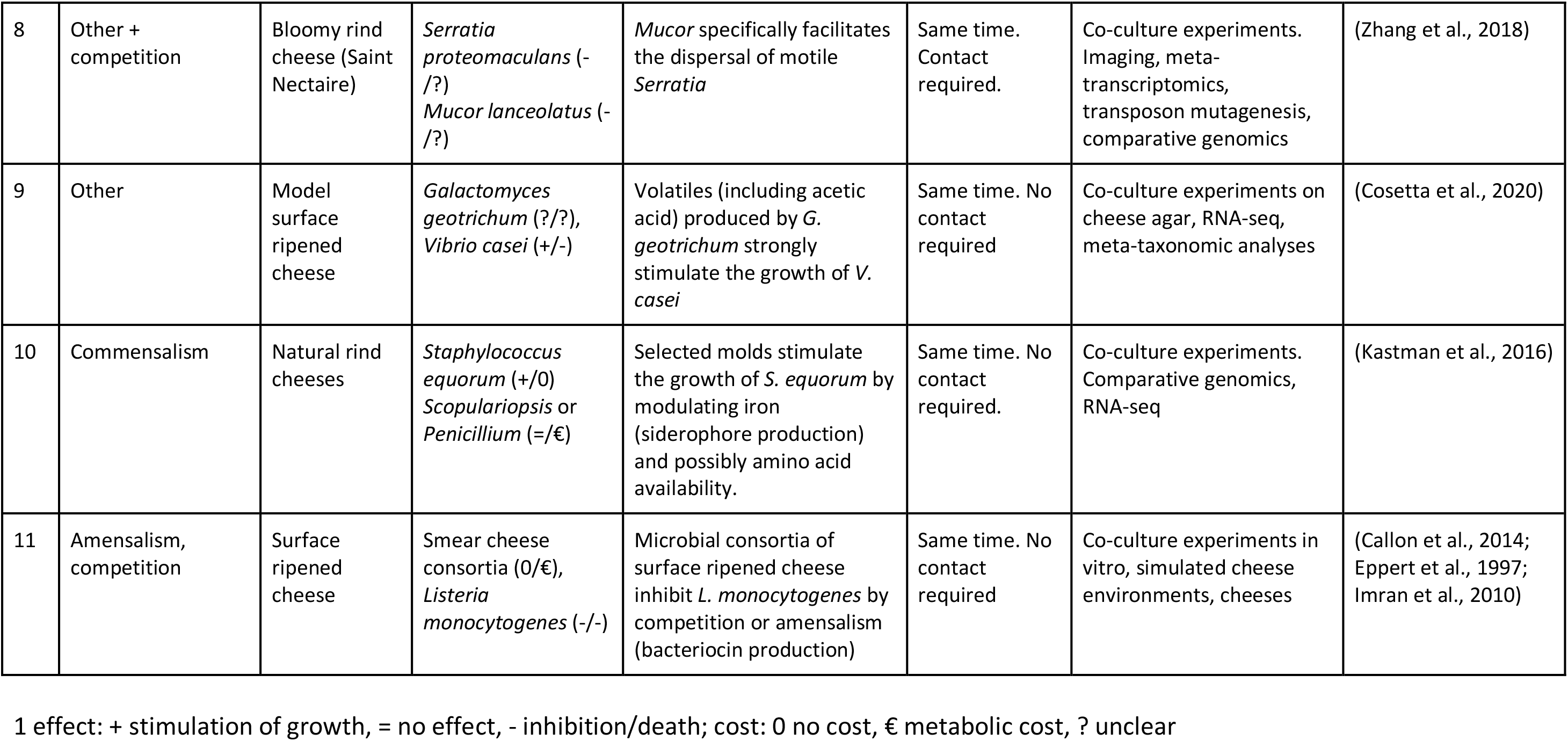
Some examples of microbial interactions between cheese microorganisms and of methods used for their study in model systems or in cheese

Microbial interactions in cheese are best described using network concepts. Surprisingly, while network science approaches are frequent in the study of host and environmental microbiomes, they are much less so in food and dairy microbiology (Parente et al., 2018). This is rather unfortunate, given the potential of the study of microbial interactions in food for the development of new processes and products and for the optimization of microbiome intervention strategies even in complex communities (Canon et al., 2020).

In this review, we will briefly illustrate network approaches to the study of microbial communities, related terminology and methods. We will then review recent literature using high-throughput sequencing or meta-omic approaches for the study of microbial interactions in cheese microbial communities. Finally, we will exploit the metataxonomic data stored in the DairyFMBN database (Parente et al., 2020) to provide examples of the mining of microbial associations in cheese.

## 2. Network analysis concepts and approaches to the study of microbiota

### 2.1 Network science concepts

A **network** is, simply put, a collection of objects (**nodes**, vertices) connected by interactions (**edges**, arcs, links) (Newman, 2010). Because of the flexibility and power of this concept, network science is used in representing and understanding interactions in very diverse fields (physics, social sciences, biology, etc.). In the study of microbial associations, nodes can be of the same type (**unipartite networks**, as in microbial association networks, in which the nodes are microbial taxa, Operational Taxonomic Units, OTU, or Amplicon Sequence Variants, ASV) and the edges represent some sort of true or inferred (positive or negative) interaction. However, **bipartite networks** (i.e. networks with nodes belonging to two different types) are also of interest: networks of this type include Phage-Bacteria Interaction Networks (PBINs; Flores et al., 2011) and Food-Microbe interaction networks (which have been frequently used as a descriptive tool for the structure of microbial communities in foods (Parente et al., 2018). In this case, edges represent simply the occurrence of a relationship (a bacteriophage infecting a bacterial strain, the presence of a given taxon in a given sample), which can be sometimes weighted (see below). More complex networks (sample-OTU-metabolite) can be built using multi-omic data (Liu et al., 2020).

In most cases, networks used for representing microbial associations are **undirected** (i.e. the edge does not point from one of the nodes to another, because the relationship is considered to be reciprocal). On the other hand, parasite-host networks are usually represented as **directed** networks, with edges going from phages to bacteria. Directed networks can be used to represent in a more meaningful way ecological relationships like commensalism, amensalism or parasitism (while mutualistic relationships and competition are typically undirected) (Canon et al., 2020; D’Souza et al., 2018).

Networks can be **weighted** if the edges have some sort of integer or real number associated to them, representing the “strength” of the association (i.e. a correlation or distance value for association networks, abundance of a taxon in a given sample, some measure of the virulence of a phage towards a host, etc.).

Networks can be characterized by many network- and node- or edge- level indices. A detailed review of these indices is beyond the scope of this work, and interested readers are referred to comprehensive reviews on microbial interaction networks (Layeghifard et al., 2017; Liu et al., 2020). Besides the number of nodes and edges, the **average degree** (the average number of edges per node), the **connectance** (i.e. the ratio between the actual number of interactions with the potential number of interactions (Delmas et al., 2019; Dunne et al., 2002), the **average path length** (the average distance between pair of nodes), the **node degree distribution** (the distribution of the probability that a node has a given degree) and the **global clustering coefficient** (a measure of the organization of the network in modules or cliques with a higher average degree among them than with other nodes; this can also be measured by **modularity**) are all important measures of network structure. The latter properties, taken together, allow to typify network topologies (Layeghifard et al., 2017). Microbial association networks in host and environmental biomes have been found to differ significantly from random networks (i.e. networks in which each node has an equal probability of having an edge with another node, and in which the node degree distribution follows a Poisson distribution) and to have a **scale-free** structure (with a power law node degree distribution, in which most nodes have a small degree while some, the hubs, have a large number of edges) with **small-world** properties (low average path length, highly modular structure). The occurrence of highly interconnected hubs and of densely connected modules of nodes should result in resistance to disturbance (i.e. removal of random nodes or edges should not significantly change the structure of the network). On the other hand, removal of hub species or of edges connecting different modules can result in significant disruptions. On the other hand, food MAN have been found to be simpler, to lack the small-world, scale free structure of environmental MAN, and to have at most a truncated power-law distribution of node degrees (Layeghifard et al., 2017; Parente et al., 2018; Röttjers and Faust, 2018). Since MAN can include both positive and negative associations, another important network property is the **proportion of positive edges (PEP)** (Faust et al., 2015). Changes in the ratio of positive to negative interactions may be considered markers for shift from a “healthy” to a “diseased” microbiome (Ma, 2018).

In general, comparison of sets of global network properties may help in understanding the global impact of changes in a food microbiome due to intervention, spoilage or fermentation (Parente et al., 2018; Peschel et al., 2020).

Finally, several individual node properties (**degree, weighted degree, other centrality measures, clustering coefficient**) contribute to the identification of keystone species in microbial association networks (Layeghifard et al., 2017; Liu et al., 2020; Röttjers and Faust, 2018). **Hubs** are characterized by high values of centrality measures (degree, closeness and eigenvector centrality) while **bottlenecks** are characterized by high values of betweenness centrality (the number of shortest paths passing through a node) and both may have key roles in ecosystem functioning and stability. Finally, the detection of modules of highly interconnected nodes may help in the identification of groups of taxa which share the same niche or may have strong metabolic interactions. It is worth noting that betweenness can be also calculated for edges: edges which have high **edge betweenness** are important for the structure of the network, because their removal disrupts the network.

### 2.2 Methods for the inference and visualisation of microbial association networks

Because of the importance of network approaches in the description and understanding of microbial communities, a large variety of methods have been developed for the representation and inference of microbial association networks.

Visual representation and analysis of bipartite sample-OTU networks (in which edges simply represent the occurrence of a OTU in a sample, possibly weighted by the relative abundance) has and is been used as a descriptive tool for the identification of clusters of samples or taxa and for the identification of the core and accessory microbiota (Parente et al., 2016). Node and edge tables derived from metataxonomic data are frequently imported in graph visualisation software (like Gephi, https://gephi.org/, or Cytoscape https://cytoscape.org/; Shannon et al., 2003). These tools allow to filter the nodes, calculate network and node statistics, use them or node or edge metadata to apply styles (shapes, sizes, colors, etc.) to subsets of nodes and edges, and to rearrange the graph using layout algorithms with the purpose of identifying modules or hubs.

The inference of microbial associations (or of more complex associations in meta-omic data) requires specialized tools (Liu et al., 2020; Peschel et al., 2020; Röttjers and Faust, 2018). Unfortunately, interactions between taxa, or between taxa and metabolites, are still being inferred from correlation (either Pearsons’s r or rank order correlations) matrices using some sort of arbitrary threshold for sparsification (i.e. removal of edges which fall below a given correlation threshold), or worse still, some sort of testing of the “significance” of the correlations (Liu et al., 2020; Peschel et al., 2020; Röttjers and Faust, 2018). The most frequently used data are 16S rRNA marker gene data, which are affected by **sparsity** (a large number of zeros, for which it is usually difficult to establish if they are simply missing data due to insufficient sequence depth or if they are structural zeroes, reflecting the absence of a given feature) and **compositionality** (the true abundance of the target gene is rarely measured and therefore relative frequencies are generally used; even when absolute abundances are used, they suffer from a compositional bias, since, when the increase in abundance of a given taxon must be accompanied by the decrease in abundance of less abundant taxa). In addition, in several cases, **indirect correlations** may be detected (i.e. an association between taxa A and B is inferred if both are associated with taxon C or with a common environmental condition). Further difficulties ensue from the limitations of amplicon targeted approaches, whose resolution is often limited to the genus level. Finally, time-series data with enough time points to use model based approaches for the inference of microbial interactions are relatively rare (Röttjers and Faust, 2018), and methods developed for cross-sectional data are more frequent.

A complete review of the methods and algorithms used to infer microbial association networks for cross-sectional data, of their strengths and weaknesses is beyond the scope of this paper: this is an active area of research and scores of papers have been published in recent years. The interested readers are encouraged to peruse one of the many recent reviews on the subject (Jiang et al., 2019; Liu et al., 2020; Röttjers and Faust, 2018). A few approaches have been used more frequently or tested in comparative studies and are available in NetCoMi, a R package specialized in microbial association network inference and network comparisons (Peschel et al., 2020). **SparCC (Sparse Correlations for Compositional data**; Friedman and Alm, 2012) infers networks based on Pearson correlations on log-ratio transformed data, thus addressing, at least in part, the issue of compositionality, and has been frequently used. **CCREPE** (Compositionality Corrected by REnormalization and Permutation; Faust et al., 2012), also known as ReBoot, uses permutations and renormalization to remove correlations due to compositionality alone and is implemented in the CoNet app (Faust and Raes, 2016) with a variety of correlation, similarity and dissimilarity based measures, which can be used in an ensemble approach. **SPIEC-EASI** (SParse InversE Covariance Estimation for Ecological Association Inference; Kurtz et al., 2015) is based on estimation of conditional covariances, is robust to both compositionality and indirect correlations, and has performed well in benchmarking, especially when using the neighborhood selection method (also known as the MB method (Meinshausen and Bühlmann, 2006). Another recent method based on estimation of semi-parametric correlation and inference of conditional dependence is **SPRING** (SemiParametric Rank-based approach for INference in Graphical model; Yoon et al., 2019).

More recently, high throughput methods for large scale network inference which are also able to remove indirect correlations by including information on sample properties have been made available (Tackmann et al., 2019).

Network inference is method specific, and specific interactions, like amensalism, may be undetectable for cross-sectional studies (Weiss et al., 2016), but we have recently shown that combining several methods may help in detecting stable and scientifically reasonable associations in food microbiomes (Parente et al., 2018).

The scarcity of studies on the inference of microbial association networks in cheese (and in food in general) may have been, at least in part, due to the lack of user-friendly software for network inference, characterization and comparison, and, possibly, to the lack of large publicly available databases. This has changed recently, as comprehensive and relatively user-friendly R packages have become available (Peschel et al., 2020). Web based platforms, like MetagenoNets (Nagpal et al., 2020) have also become available, thus empowering even users lacking coding abilities. While a naïve use of these tools should be discouraged, their availability, in conjunction with complex metataxonomic databases as FoodMicrobionet and DairyFMBN (Parente et al., 2019, 2020) should significantly facilitate the mining of existing and new data for microbial associations.

### 2.3 Confirming interactions in microbial association networks

Computational tools for the inference of microbial association networks are essentially explorative tools. They can be used to provide hints for the existence of novel patterns of interactions, but processes and mechanisms underlying the interaction must be confirmed by *in-vitro* or *in-vivo* experiments (Cosetta and Wolfe, 2019). Interactions in cheese are frequently mediated by diffusible or volatile metabolites (including volatile organic compounds) and/or by physical contact.

With the exception of experiments performed *in-vivo* (i.e. in real cheese systems), simplified communities (2-4 members) are most frequently used, even if much larger assemblages have been tested in some cases (Callon et al., 2011, 2014; Imran et al., 2010). Simplified communities grown in liquid media are clearly the easiest way to clarify the mechanisms of microbial interactions, but they may be insufficient to resolve the complex and dynamic interplay in cheese, especially in cheeses produced with undefined starters, raw milk varieties or surface ripened cheeses.

In fact, several commensalistic interactions (SLAB-NSLAB in most cheeses ripened internally by bacteria; LAB-propionibacteria in Swiss-type cheeses; LAB - fungi - surface flora in surface ripened cheeses; Blaya et al., 2017; Gobbetti et al., 2018; Mayo et al., 2021; Sieuwerts et al., 2008; Smid and Lacroix, 2013) are, in fact, indirect, and due to niche conditioning (change of pH, production/consumption of substrates, release of nutrients) and most of the growth of the partners occurs in different life stages of the cheese (curd manufacturing, ripening) or at different locations (core, surface). The interested reader is referred to the many excellent reviews which have been published on this subject. Here, we will concentrate on more direct interactions.

Table 1 summarizes some representative interactions, together with the experimental approaches used for their study, ranging from simple growth experiments, occasionally with modelling of growth kinetics, to meta-transcriptomic and metabolomic approaches in cheese or model systems. Most studies analyze commensalistic relationships, either due to ecosystem conditioning or to exchange of metabolites, and the majority is focused on surface ripened cheeses and bloomy rind cheeses, in which the development of desired sensory properties relies on complex interactions between lactic acid bacteria, *Proteobacteria*, *Actinobacteria*, *Staphylococcaceae*, yeasts and molds.

Negative interactions due to bacteriophages (parasitism) are relatively simple to study and have a profound impact on the structure and dynamics of microbial communities in cheese (Erkus et al., 2013; Pujato et al., 2018; Zotta et al., 2021). Because of their importance, bacteriophage - host interaction networks are reviewed in a separate paragraph.

Amensalism due to production of bacteriocins is also important (both because it might affect the stability of mixed strain starters or because of potential uses in bio-preservation and control of pathogenic and spoilage organisms; Silva et al., 2018), and relatively easy to study in vitro (see Favaro et al., 2015; Lozo et al., 2021 for recent reviews). In addition, mining metagenomes for bacteriocin genes (Escobar-Zepeda et al., 2016; Lozo et al., 2021; Walsh et al., 2020), and using targeted methods for studying their expression in cheese is relatively easy (Trmčić et al., 2011) but, to our knowledge, there is no meta-transcriptomic data on the expression of bacteriocin genes in cheese during ripening.

Complex interactions are responsible of anti-listerial activity of microbial consortia which develop on surface-ripened cheeses. The anti-listerial activity may be due to factors other than bacteriocin production, including competition, and studies in model systems (cheese agar) have shown that complex consortia are needed, and that activity varies significantly with their composition and complexity (Callon et al., 2014; Imran et al., 2010).

Amensalism can be strongly affected by the solid nature of the cheese matrix: cheese moisture and even the internal environment of colonies may affect the diffusion of relatively large molecules (Floury et al., 2015; Guitián et al., 2019) like bacteriocins but also, in the short term, of smaller molecules.

In fact, spatial distribution of colonies and diffusion of molecules within and between colonies in or on the cheese matrix may significantly affect cheese quality for several reasons. First of all, except when cells or microcolonies are in close proximity, interactions are probably not relevant. Gradients within a single colony may be important in affecting the growth in different locations of a colony (Malakar et al., 2000, 2003), although the buffered environment of cheese may prevent the existence of pH microgradients, at least in microcolonies (Jeanson et al., 2013). Inoculum level may affect size and distribution of microcolonies and this, in turn, has been shown to affect cheese composition (Boucher et al., 2015a, 2015b; Jeanson et al., 2010), although, in the long term, it might not affect the distribution of small soluble metabolites implicated in some cross-feeding relationships SLAB and LAB, (Czárán et al., 2018) at least in internally ripened cheeses.

The situation is quite different in mold ripened cheeses and surface ripened cheeses. In fact, the composition of the microbiota on the surface and core of several cheeses (including cheese internally ripened with bacteria) is significantly different (see Jonnala et al., 2018 for a review) and differences in ecological conditions may strongly affect interactions between microorganisms and vice-versa. In Stilton cheese, the occurrence of mixed micro-colonies close to the internal veins formed by the mold was shown by Fluorescence In Situ Hybridization (Ercolini et al., 2003) and this was hypothesized to be due to commensalism. However, in the same cheese, lactococci and leuconostoc formed in other parts of the curd pure culture microcolonies, and microcolonies identified as *Lactiplantibacillus plantarum* or tentatively identified as *Latilactibacillus curvatus* appeared in different location (underneath the crust or close to the veins).

In conclusion, imaging techniques are certainly most useful in studying the spatial organization of interacting microorganisms in cheese (Hickey et al., 2015), but mass-spectrometry techniques may also be of great importance in evaluating the importance of compounds like siderophores (Cleary et al., 2018) or possibly bacteriocins (Hindré et al., 2003).

Microbial interactions on cheese rinds may affect in a complex way the survival and growth of *Listeria monocytogenes* and *Escherichia coli*, two pathogens which have been associated with foodborne outbreaks due to the consumption of surface-ripened cheeses (Fusco et al., 2020).

Co-cultivation with bacteria found on the surface of cheese (*Brevibacterium*, *Psychrobacter*) has been found to affect the expression of genes whose transcription is regulated by the global stress regulator σ_B_ (Anast and Schmitz-Esser, 2020) and the expression of a small non coding RNA, thus suggesting that stress due to competition is involved in this interaction. Random Barcode Transposon Sequencing and RNA-Seq have recently been used to identify genes which are related to the survival and growth in a cheese model of *Escherichia coli*, when grown alone or in binary and multiple associations with cheese-rind microorganisms like *Geotrichum candidum, Hafnia alvei* and *Penicillium camemberti* (Morin et al., 2018). Important genes were related to amino acid synthesis or metabolism, iron acquisition of response to toxic compounds and oxidative stress, thus confirming one more the importance of these factors in fitness and interactions in cheese. In addition, growth in co-culture demonstrated that the cheese species might exert both positive effects (by providing free amino acids) and negative effects (by amensalism through the production of toxic compounds or oxidative stress). Incidentally, this experiment also demonstrated that pairwise interactions may be not representative of the growth in more complex (4 members) communities. Although this highlights the limitations of experiments in model systems, teasing out interactions in real systems may prove exceedingly difficult, due to variability in time and space and to the complex and dynamic nature of the microbial communities, especially in surface ripened cheeses.

The collection and interpretation of multi-omic data is probably the most promising approach for the study of microbial interactions in cheese during curd production and ripening, although cost and computational power issues may still limit the collection of large longitudinal data sets needed to use model based approaches for inferring interactions. The most extensive demonstration of the value of complex, integrated approaches, for finely dissecting the complex interactions in cheese microbial communities is probably the on-going work on surface ripened cheeses (Bonham et al., 2017; Cleary et al., 2018; Cosetta and Wolfe, 2020; Cosetta et al., 2020; Kastman et al., 2016; Niccum et al., 2020; Wolfe et al., 2014; Zhang et al., 2018). Starting from an extensive metataxonomic and metagenomic characterization of the rind microbial communities of bloomy surface, washed rind and natural dry rind (Wolfe et al., 2014) from all over the world, and from the development of a detailed set of protocols for dissecting interactions in model cheeses (Cosetta and Wolfe, 2020), this group was able to provide detailed evidence for the drivers which determine the diversity and functionality of microbial communities on the surface of cheeses.

In fact, surface ripened cheeses provide an excellent model for the study of microbial interactions in an environment which, at least in part, can be mimicked in the laboratory under the controlled conditions which are needed to dissect the nature of the interactions and establish cause-effect relationships.

First of all, cheeses which are surface ripened by fungi (thus resulting in a bloomy rind, like Camembert, Brie, Saint Nectaire; Spinnler, 2017) or by complex consortia of yeasts, molds and bacteria (washed rind cheeses, like Limburger, Tilsit, Port du Salut, Taleggio, etc.; Mounier et al., 2017) are economically important worldwide and include both traditional varieties (which are still produced by using raw milk and rely on natural contamination from the environment, with very limited use of starter cultures) and industrial varieties, which are produced using pasteurized milk inoculated with defined or undefined starters and in which the development of the surface microbiota is promoted by the addition of specific combinations of fungi and bacteria. The composition of the rind microbiota of these cheeses is significantly more complex that those of cheeses with hard, dry rind (Wolfe et al., 2014), and the mature microbiota is the result of a complex succession: growth of halophilic, acid sensitive bacteria is only made possible by the consumption of lactic acid by yeasts and by the increase in pH due to proteolysis caused by molds. Recent research using high-throughput sequencing approaches has shown that, beyond the complex assemblages of *Actinobacteria* (*Brevibacterium, Microbacterium, Arthrobacter, Corynebacterium*) and *Firmicutes* (*Staphylococcus*) which have been traditionally been associated with the pigmentation and aroma of these varieties, several *Proteobacteria*, including *Pseudoalteromonas, Hafnia, Vibrio, Halomonas,* and *Psychrobacter* may contribute with their metabolic activities (Afshari et al., 2018; Jonnala et al., 2018; Wolfe et al., 2014). In addition, the increase in pH of the rind during ripening makes the growth and survival of pathogenic microorganisms easier, including *Listeria monocytogenes* and Shiga-toxin producing *Escherichia coli*. Understanding how these species are controlled by amensalism and competition is a key factor for the safety of these cheeses. Finally, since several subdominant species which are prevalent on the surface of these cheeses are not members of the starter cultures, the role of cheesemaking environments, including different areas in the cheese plant and ripening shelves in the dispersal of these species and in the maintenance of a continuous inoculation source is also important in both providing beneficial and spoilage microbes (Bokulich and Mills, 2013; Guzzon et al., 2017).

Fungi (both yeasts and molds) and bacteria coexist on the surface of several cheeses and, although environmental and technological conditions may determine some of the co-occurrence and co-exclusion patterns, microbial interactions (both trophic and non trophic) do explain why some of these relationships systematically occur across cheeses from different geographical areas (Wolfe et al., 2014). Apart from indirect interactions (de-acidification of the cheese surface due to growth of yeasts or molds), several direct interactions are known to exist between bacteria and fungi (Mayo et al., 2021; Mounier et al., 2008). Competition for microelements, with interactions mediated by exchange of siderophores or other metal chelators is frequent in surface ripened cheeses as is competition for folic acid, which can be alleviated by the exchange of corrinoids, a frequent mechanism for microbial interactions (see Table 1; Abreu and Taga, 2016). Below, two examples from recent studies are described in more detail.

Competitive relationships may hide more subtle relationships between partners: on the surface of bloomy rind cheeses motile proteobacteria like *Serratia proteomaculans* can exploit the network of hyphae formed by fungi like *Mucor lanceolatus* for dispersal: motile cells move along the water film on the lax hyphal growth formed by the fungus to disperse on the surface of the cheese, thus colonizing a larger area (Pujato et al., 2018). The relationship is somewhat specific and no dispersal or significantly lower dispersal is observed with other cheese fungi forming denser hyphal networks. On the other hand, as in many other cases, some competition is observed between the two partners, because both show less growth when grown together than when grown alone. Dispersal ability also varies among strains of *Serratia*. RNA-seq analysis of *Serratia* grown in co-culture with *Mucor* showed that co-growth alters the supply of nutrients and metabolites and the metabolism of *Serratia*, but no significant difference in the expression of genes related to motility and quorum-sensing was found. Both transposon mutagenesis and comparative genomics supported the fact that genes related to flagellin production (and hence to motility) are essential for dispersal, but also provided proof that other genes are implicated in the relationship between these two species: a few mutants were able to kill *Mucor*, while others showed similar dispersal but altered colony morphology. In the same paper the authors elegantly demonstrated that fungal networks dramatically affect the structure of the microbial community and that dispersal, as well as other interactions (competition, amensalism) are implicated. In fact, several motile *Proteobacteria* could disperse on fungal networks, while motile and non motile species of *Firmicutes* and *Actinobacteria* showed limited dispersal or no dispersal. Fungal growth may be therefore responsible for better growth of *Proteobacteria* on the surface of bloomy rind cheeses. Growing hyphae can be, at least in part, replaced by physical, inert and non-growing networks of glass fibers.

While many interactions may require a more or less close contact between partners, some are mediated by volatile compounds. Recently (Cosetta et al., 2020) it has been shown that volatile compounds produced by molds isolated from bloomy or washed rind cheeses (*Galactomyces geotrichum*, *Debaryomyces hansenii*, *Penicillium* sp., *Scopulariopsis* sp., *Fusarium domesticum*) strongly stimulate or inhibit the growth of selected *Proteobacteria* (with *Vibrio* being strongly stimulated, and *Pseudomonas* often inhibited), and contribute to explaining the abundance of these species on cheese rinds. In the case of *Vibrio casei*, stimulation was related to significant changes in the transcriptome, and free fatty acids and esters of free fatty acids produced by the fungal partners were implicated in the relationship.

## 3. Bacterial association networks in cheese. A meta-study

The inference of microbial association networks is surprisingly rare in studies dealing with the structure and dynamics of cheese microbiota (Murugesan et al., 2018; Parente et al., 2018), although simple bipartite network representations (De Filippis et al., 2014; Guidone et al., 2016) have sometimes been used with the main objective of visually representing the core microbiota. In a previous paper (Murugesan et al., 2018; Parente et al., 2018) we have inferred microbial association networks for 8 cheese metataxonomic datasets extracted from FoodMicrobionet using three inference methods (CoNet, SPIEC-EASI and SparCC) and compared them with those from other food, host or environmental biomes. In the meantime, the size of FoodMicrobionet has grown substantially (Parente et al., 2019) and a specialized version for dairy foods (DairyFMBN; Parente et al., 2020) has been made available. Version 2.1.6 of DairyFMBN (https://data.mendeley.com/datasets/3cwf729p34/5) includes 74 studies and 4,738 samples. This offers the opportunity to mine a large set of data, which albeit heterogeneous (DairyFMBN includes metataxonomic studies with different designs and gene targets) is highly structured, and to obtain insights on the approaches which can be best used to infer association networks and, possibly, on conserved interactions and keystone species which may warrant further study.

We therefore extracted 34 studies which had at least 20 samples (Supplementary Table 1) and processed them using a pipeline based on NetCoMi (Peschel et al., 2020). An example workflow is available from GitHub (https://github.com/ep142/MAN_in_cheese) and the details of the approach are described in Supplementary material. The workflow allowed rapid and reproducible analysis of small to medium sized datasets and was used to answer different methodological questions.

### 3.1 Which is the most appropriate taxonomic level for the inference of association networks?

This is probably the first and most important challenge in a study aimed at inferring microbial interactions. The lowest possible taxonomic level would, of course, be desirable, because it has been proven that microbial interactions in cheese can be strain specific (Mayo et al., 2021; Niccum et al., 2020). On the other hand, even if Amplicon Sequence Variants may be preferable to OTUs and their inference may offer some insight in diversity below the species level (Callahan et al., 2016, 2017; Porcellato et al., 2021), and even if, with proper care and with the use of dedicated databases, identification at the species level can be significantly improved (Meola et al., 2019; Pollock et al., 2018) heterogeneity in wet- and dry laboratory protocols may make comparison between different studies possible only at higher taxonomic levels (genus; De Filippis et al., 2018; Pollock et al., 2018). In our dataset, 62% of the studies were processed using a pipeline based on DADA2 (Callahan et al., 2016, 2017; Parente et al., 2019; Porcellato et al., 2021) (from study 34 to 165, Supplementary Table 1) but the target and the sequence lengths were variable (Supplementary Table 1). In fact, the median value of the proportion of sequences (within study) identified at the species level was 0.176 (range 2×10^−5^−0.99), while the median value of the proportion of sequences identified at the genus level or below was 0.996 (range 0.25-1). Agglomeration at the genus level might be a necessity if one wants to compare networks from different studies and it has even been claimed that, when species belonging to the same phylogenetic group share similar functions, agglomeration can reduce noise (Röttjers and Faust, 2018). This might not be true for many microbes in dairy products, because, for example, genera like *Streptococcus*, *Staphylococcus* and *Corynebacterium* include both beneficial and pathogenic species. To illustrate this point we carried out inference of microbial association with four methods (see the description of the workflow in Supplementary material), two based on correlations (SparCC and CCREPE) and two on conditional dependence (SPRING and SPIEC-EASI), on datasets in which sequences were not aggregated (Amplicon Sequence Variants, ASVs, prior to import in DairyFMBN), or with no taxonomic aggregation (ASVs were pooled at the lowest possible level of taxonomic assignment) or with aggregation at the genus level. The inferred networks with CCREPE and SPIEC-EASI for DairyFMBN data with no taxonomic aggregation for two V3-V4 targeted studies (ST49, a large study on the microbiota of a diverse set of artisanal cheeses from Brazil, (Kamimura et al., 2019) and ST136, a study on the impact of starter addition on microbiota of fromadzo cheese, (Dolci et al., 2020); because of sequence quality taxonomic identification for ST49 was at the genus level or above) producing relatively simple networks are shown as an example in Figure 2. The level of identification of taxa varied from species (27% and 17% of the sequences identified at the species level for ST49 and ST136, respectively), to family (poorly identified taxa were removed by filtering), while networks inferred using ASVs or taxa aggregated at the genus level are shown in Supplementary Figure 2 and 3, respectively.

**Figure 2.**
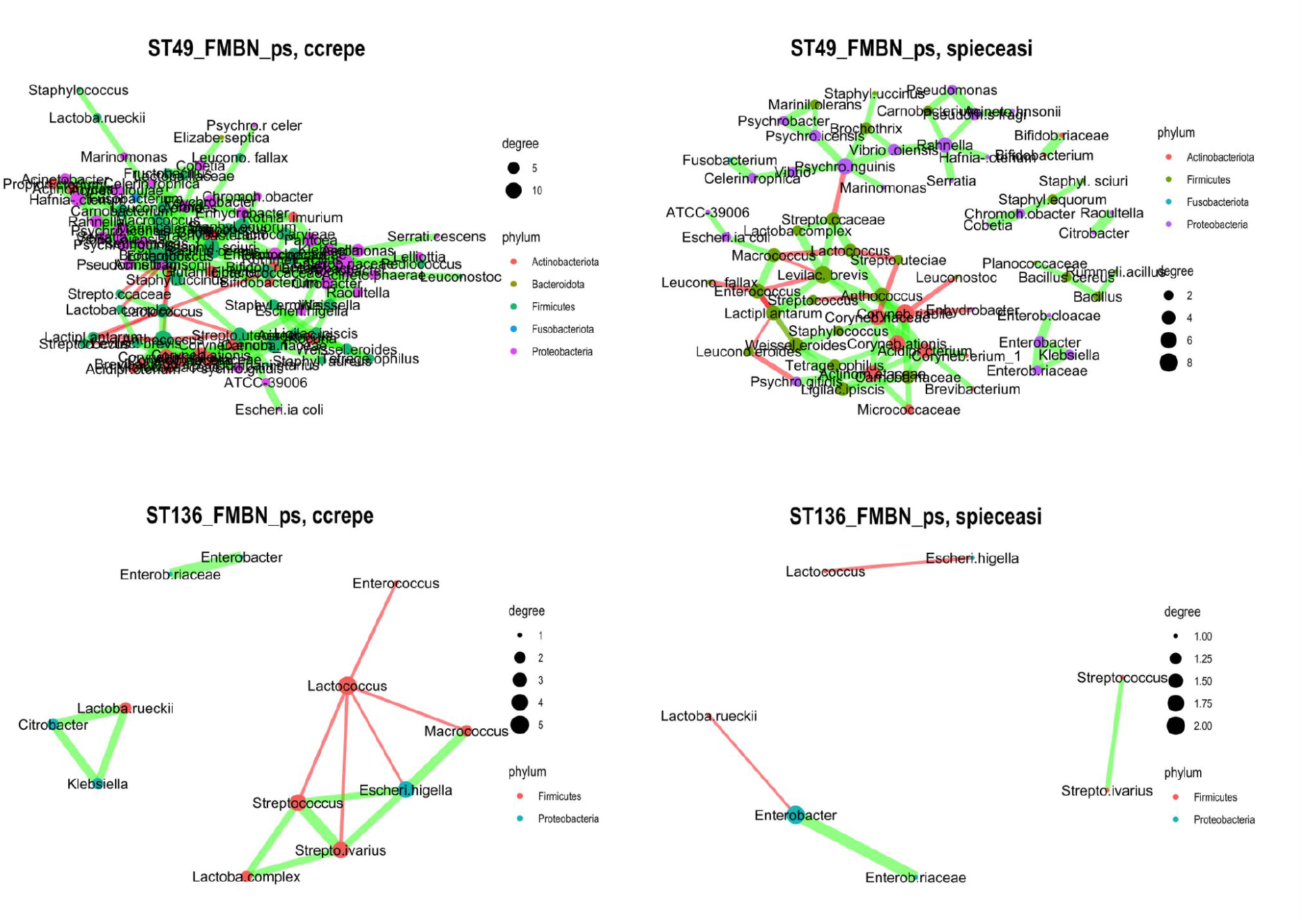
Microbial association networks inferred from studies ST49 (Kamimura et al., 2019) and ST136 (Dolci et al., 2020) extracted from DairyFMBN 2.1.6, with no taxonomic aggregation (https://data.mendeley.com/datasets/3cwf729p34/5) using methods CCREPE (Faust et al., 2012) or SPIEC-EASI (Kurtz et al., 2015). For each pane, the color of nodes is determined by phylum, and the size by degree; the color of the edges is red for mutual exclusion relationships and green for copresence relationships; the thickness of the edges is determined by the strength of the association measure, as determined in NetCoMi (Peschel et al., 2020). A force-based layout (Fruchterman–Reingold) was used for positioning nodes and edges. The name of the nodes has been abbreviated to 15 characters.

Networks based on ASVs (Supplementary Figure 1) and those obtained from DairyFMBN data without taxonomic agglomeration have, as expected, the largest number of nodes and include nodes identified at the species level. The interpretation of the networks is beyond the scope of this meta-study but several negative interactions between starter bacteria and spoilage species or positive interaction between potentially beneficial species or between spoilage species can be identified. It is also evident that, at this level of aggregation, parent-child relationships (i.e. a positive association between nodes belonging to the same genus, one of which is identified at the species or genus level, while the other is identified at a higher taxonomic level; i.e. *Streptococcus* -- *Streptococcus salivarius* or *Enterobacter* -- *Enterobacteriaceae*) are evident. In fact, the median value for the proportion of parent-child relationships ranged from 0.03 for networks inferred by CCREPE to 0.067 for networks inferred using SPIEC-EASI, but was as high as 1 for some very sparse networks. This is even more true for the networks based on ASVs, in which clusters of nodes belonging to the same genus and connected by copresence relationships are clearly evident. Although ASVs offer a higher resolution than nodes obtained from taxonomic agglomeration, their biological meaning is not clear and microbial association networks tend to have so many nodes and edges that they are hardly interpretable (although node filtering on the basis of some centrality measure can be performed). On one hand it is very well known that most bacterial species have multiple copies of the 16S rRNA genes which differ in their sequence (Větrovský and Baldrian, 2013) and, although it might be questionable that partial sequencing may be sufficient to distinguish variants within and between strains (Johnson et al., 2019) it is not uncommon that more than one ASV is detected for each strain in mock communities (we detected on average 2 ASVs per microbial species in ZymoBIOMICS™ Microbial Community DNA Standard, data not shown), but it is unclear if this is due to sequencing errors or to the detection of different copies of the 16S rRNA gene.

Likewise, ASVs belonging to a cluster of nodes with the same genus or species identification may, in fact, belong to different co-occurring strains or simply be the result of sequencing errors which may lead to identification at different taxonomic levels. The effect of parent-child relationships or of copresence edges between nodes belonging to the same higher taxon (ASVs belonging to the same species, species belonging to the same genus) is clearly seen by comparing the plots for ASVs, species and genus, with cluster of taxonomically related nodes progressively collapse in a single node.

The choice of the taxonomic resolution level is therefore related to the quality and resolution of the data and to the objective of the study. Using ASVs is probably not justified. Whenever the quality and length of the sequences allow it, the aggregation at the lowest possible taxonomic level should be used (this is not the same as using taxonomic agglomeration at the species level in phyloseq, because this would remove all ASVs identified above the species level). However, when comparing different studies aggregation, the genus level is probably desirable, although this may blur or hide some associations at lower taxonomic levels.

### 3.2 Which is the best inference method?

It is well known that inference methods for microbial associations differ in their ability to detect copresence and mutual exclusion associations, and that the inferred networks are method specific and may be affected by the inclusion of environmental variables in the inference (Tackmann et al., 2019). This has been reviewed multiple times (Liu et al., 2020; Röttjers and Faust, 2018; Weiss et al., 2016) and further discussion is beyond the scope of this paper. However, the availability of tools which allow rapid inference of networks with multiple methods (such as NetCoMi; Peschel et al., 2020) makes this task relatively easy (an example workflow is available from GitHub: https://github.com/ep142/MAN_in_cheese). We therefore carried out network inference on 34 cheese datasets included in DairyFMBN (see Supplementary Table 1) using 2 correlation based methods (SparCC and CCREPE) and two conditional dependence methods (SPRING and SPIEC-EASI) (see section 1.2). All methods are able to handle compositionality well (although they use different methods), and their requirements in terms of number of samples and taxa or in terms of sparsity of the underlying network differ. SparCC and CCREPE are included mostly for historical reasons while SPIEC-EASI has repeatedly showed good performances in network inference compared to other methods (Kurtz et al., 2015; Tackmann et al., 2019; Weiss et al., 2016; Yoon et al., 2019). SPRING is a relatively novel method, which has been claimed to offer superior performances (Yoon et al., 2019) but which, to our knowledge, has not been tested on food microbiomes.

A comparison of two methods for two studies was shown before see 3.1). Figure 3 shows a Venn diagrams of edges for two further studies: ST41, a V3-V4 targeted cross-sectional study on French surface ripened cheeses (Dugat-Bony et al., 2016) and ST131, a V4 targeted time-series study on Cheddar (Choi et al., 2020), while the corresponding networks are shown in Supplementary Figures 4 and 5.

**Figure 3.**
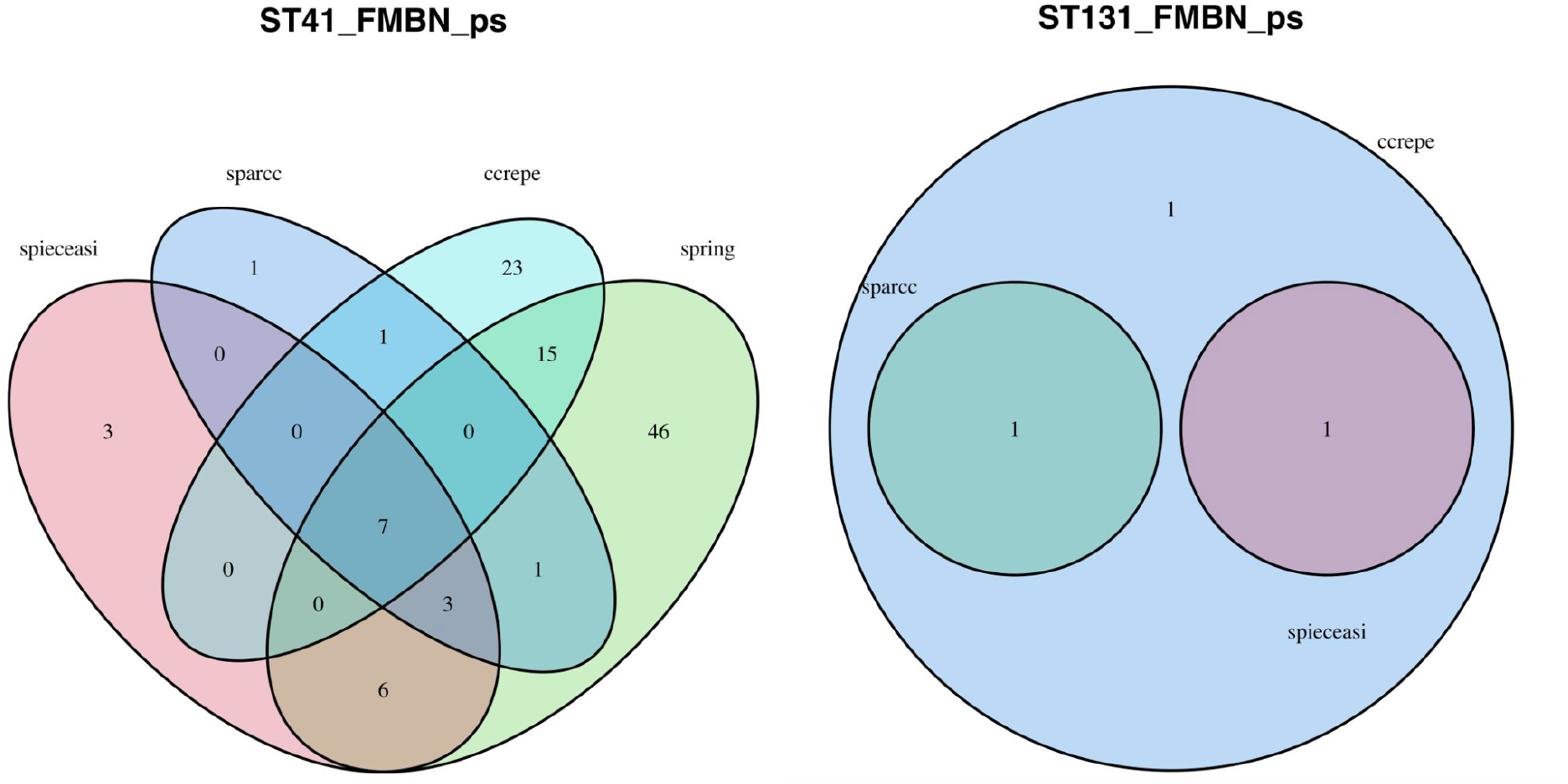
Venn diagrams for microbial association edges for networks inferred for two studies (ST41, a V3-V4 targeted cross-sectional study on French surface ripened cheeses (Dugat-Bony et al., 2016) and ST131, a V4 targeted time-series study on Cheddar (Choi et al., 2020) extracted from DairyFMBN 2.1.6 (https://data.mendeley.com/datasets/3cwf729p34/5). Network inference was carried out using four methods (SparCC, Sparse Correlations for Compositional data, Friedman and Alm, 2012; CCREPE Compositionality Corrected by REnormalization and PErmutation, Faust et al., 2012; SPIEC-EASI, SParse InversE Covariance Estimation for Ecological Association Inference, Kurtz et al., 2015; SPRING, SemiParametric Rank-based approach for INference in Graphical model, Yoon et al., 2019) after taxonomic agglomeration at the genus level, as described in Supplementary material.

First of all, as expected, microbial association networks are method specific (Figure 2, 3, Supplementary Figures 2-5), and not all methods result in the inference of a network with at least one edge for a given study (Figure 3, Supplementary Figure 5). On the other hand, some associations are conserved across two or more methods. Conversely, some edges may even represent associations of opposite signs (copresence rather than mutual exclusions) in different methods (Supplementary Figure 5).

SPIEC-EASI almost always resulted in the networks with the lowest number of edges (Supplementary Figure 6), with a significant difference from all other methods (p<0.05) as assessed using a pairwise Wilcoxon rank sum test.

SPIEC-EASI is known to remove indirect associations (i.e. associations resulting from indirect correlations, see section 1.2) (Kurtz et al., 2015). SPRING is also designed to avoid indirect associations by the use of partial correlations, but it does detect a significantly higher number of edges.

Density is a measure of how connected a network is. A comparison of density of the networks for different methods is shown in Supplementary Figure 7. SPIEC-EASI resulted in networks which had a significantly higher density (p<0.001) as assessed using a pairwise Wilcoxon rank sum test: this may be related to the fact that this method resulted often in networks with only a few connected nodes.

The distribution of positive edge proportion by method is shown in Supplementary Figure 8. Although SPRING had a narrower distribution, no significant difference was found No systematic effect of study type (cross sectional, mixed or longitudinal) or size (as measured by the number of taxa detected), or target type (16S rRNA vs 16S RNA gene, target region) was evident.

The distribution of two other important network properties, global clustering coefficient and modularity, both of which are related to the organization of the microbial association network in smaller clusters of tightly interconnected nodes (which, in turn, may represent group of taxa which share similar functions or the same microhabitat) are shown in Supplementary Figures 9 and 10. The global clustering coefficient could not be calculated for many of the sparsest networks, and this measure is available only for some of the datasets. Modularity was significantly different for networks inferred by CCREPE and SPRING compared to SparCC and SPIEC-EASI, while the clustering coefficient was significantly higher for the two correlation based methods (SparCC and CCREPE). This may reflect the existence of indirect correlations.

This is the first time some global properties of microbial association networks have been compared for networks inferred for cheese microbiota using 4 methods and such a large number of studies and the results are hard to compare with other studies. The debate on the best method for inferring microbial associations from metataxonomic data is still open. On one hand, using ensembles of methods can help in detecting “stable”, robust associations, which are method independent (Yoon et al., 2019), and performing inference with multiple methods is relatively easy with the workflow we have proposed. On the other hand, only properly designed studies with a sufficient number of samples and perhaps long time series (and the use of model based approaches), and absolute, rather than relative measures of abundances of taxa and the inclusion of ecological parameters in the analysis can help in the detection of the true structure of microbial association networks (Röttjers and Faust, 2018; Tackmann et al., 2019). However, SPIEC-EASI has definitely performed well in a number of benchmarking studies and may be selected as a conservative method for microbial association inference.

Therefore, in the following sections we will use SPIEC-EASI for network comparisons and then evaluate which associations are more conserved across methods and studies.

### 3.3 Comparing global network properties

In order to evaluate and compare global network properties network, inference was performed on the 35 datasets of Supplementary Table 1 (ST10 had both data based on 16S RNA and 16S RNA gene as a target region, which were analysed separately) using method SPIEC-EASI and the pipeline described in Supplementary material. A detailed description of the procedure used to calculate global network properties and to merge them with metadata is presented in Supplementary material.

The results can only be partially compared with those of (Parente et al., 2018) because of differences in inference methods. However, we confirmed that microbial association networks for cheese, as those for many other foods, do not have a small world, scale-free nature (the fit of the degree distribution with the power law was low), but had a relatively high density (density was calculated using only connected nodes; see Supplementary Figure 7; median 0.17, Inter Quartile Range, IQR 0.96) and positive edge proportion (see Supplementary Figure 8; median 77%, IQR 55%), and a short average path length for the largest connected component (median value 0.76., maximum 3.2). Average path length non linearly increased with network size (Supplementary Figure 1). Method SPIEC-EASI resulted in relatively sparse networks, for most of which it was impossible to calculate the global clustering coefficient. On the other hand, for many of the largest studies modularity was relatively high (see Supplementary Figure 10).

To compare the studies an approach similar to that used in a previous study was used (Parente et al., 2018). The correlation between selected variables, including network properties (average path length, avPath; density, calculated on all connected nodes, density_2; modularity; natural connectivity, i.e. the average eigenvalue of the adjacency matrix, a measure of the robustness of a graph, natConnect; number of connected nodes, nnodes; positive edge proportion, pep) and properties of the datasets (average dissimilarity avDiss: average of the Bay-Curtis distance matrix; average Chao1 index; Pielou J evenness) was calculated using the Pearson product moment coefficient. A principal component analysis was then carried out. A biplot for the first two components, which explained 64% of the variance, is shown in Figure 4.

**Figure 4.**
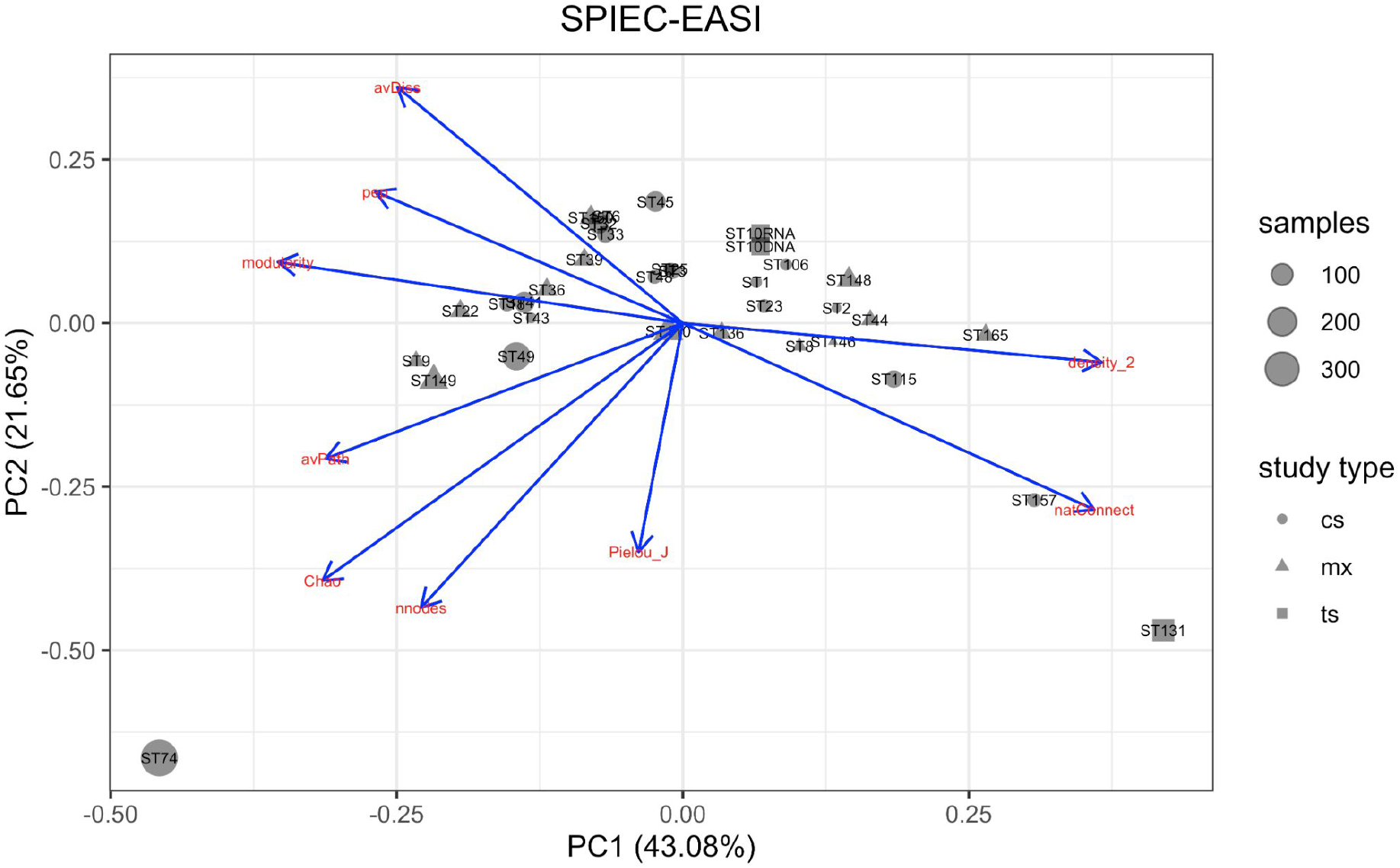
Score and loading plot for the first two components a Principal Component

Due to the heterogeneity of the studies, evaluating the results is difficult. No relationship between the study type or the target (16S RNA vs 16S RNA gene, region) is evident. The studies which stand apart are ST74, a large farm-to-fork study (Falardeau et al., 2019) which includes both environmental and milk and cheese samples, and has clearly delimited modules corresponding to different environments (feces, cheese, etc.) and a time series study on Cheddar (Choi et al., 2020) with an unusually high number of samples for each cheese/treatment and, unfortunately, a relatively low diversity due to the use of V4 region as a target and the use of pasteurized milk and defined strains starters, and ST157 (Afshari et al., 2020) which also is related to Cheddar cheese of different ages. Studies on very similar cheeses ST148, Parmigiano Reggiano (Bottari et al., 2020); ST23 Grana Padano (Bassi et al., 2015); Trentingrana (Cremonesi et al., 2020); Grana-like cheese (Alessandria et al., 2016) resulted in networks with very different properties: however, differences in study design may have obscured any similarities between the microbial communities of these cheeses.

Some aspects of alpha or beta diversity appear to be related to network properties. Average dissimilarity (as measured as average Bray-Curtis distance) was inversely correlated (−0.90) with natural connectivity, a measure of network resistance to changes, which is rather counterintuitive unless one considers than in highly modular networks the targeted removal of selected nodes connecting different clusters may result in disruption of the network. Finally, a very strong negative correlation (−0.98) was found between network density and modularity, which is expected, since high values of these properties reflect very different network structures. In conclusion, an explorative study of global network properties did not reveal any systematic pattern. Given the repeatable nature of our approach (it would be relatively easy to repeat the analysis with more data from future iterations of DairyFMBN) we feel that it should not be discarded but revaluated when more data become available.

### 3.4 Detecting keystone species

The ability of network inference approaches to detect taxa which play “special” roles in their ecosystems is one of the features which makes the analysis of microbial association networks interesting (Layeghifard et al., 2017; Röttjers and Faust, 2018), and the role of hubs (nodes with high degree or eigenvalue centrality values) or bottlenecks (nodes which may have low degree centrality but high betweenness centrality) and the effect of their removal on the stability of the ecosystem may be experimentally testable. Plots showing one or more of these properties may be useful for detecting keystone species, as shown by (Parente et al., 2018). A node plot for study ST49 (Kamimura et al., 2019) is shown as an example in Supplementary Figure 12 and similar plots can be easily obtained using our analysis workflow (https://github.com/ep142/MAN_in_cheese). In this and in other plots obtained for studies related to raw milk cheeses (data not shown) starter and non-starter lactic acid bacteria often appear as hubs with a high percentage of negative edges, while the background raw milk microbiota (whose DNA may persist during ripening) is often distributed along the dotted line which marks the average ratio between total degree and positive degree.

In principle, given a number of studies for which microbial association networks are inferred with one or more methods, one could analyse how frequently a given taxon is included in microbial association networks and evaluate how frequently it has high values for centrality values which would qualify it as a keystone. Even if we analysed a relatively large set of studies, their heterogeneity makes such an analysis difficult at this stage. However, given the repeatability of the pipeline we described, such an analysis will probably become possible in the future, as more studies are added to DairyFMBN.

### 3.5 What can edges in microbial association networks tell us?

Using an approach similar to that described above, interactions (edges) inferred for all studies using all inference methods were collected in a single data set, which can be evaluate:

a. which associations are conserved across several methods
b. given a method, which associations occur more frequently
c. which associations are important in the network structure (using edge betweenness as a measure)
d. if there is evidence of taxonomic assortativity (i.e. if the occurrence of positive associations occurs more frequently among members of the same higher taxon)

We first used the networks inferred by all four methods after taxonomic aggregation at the genus level to select for associations which were inferred by more than one method for more than one study. The results are shown in Figure 5.

**Figure 5.**
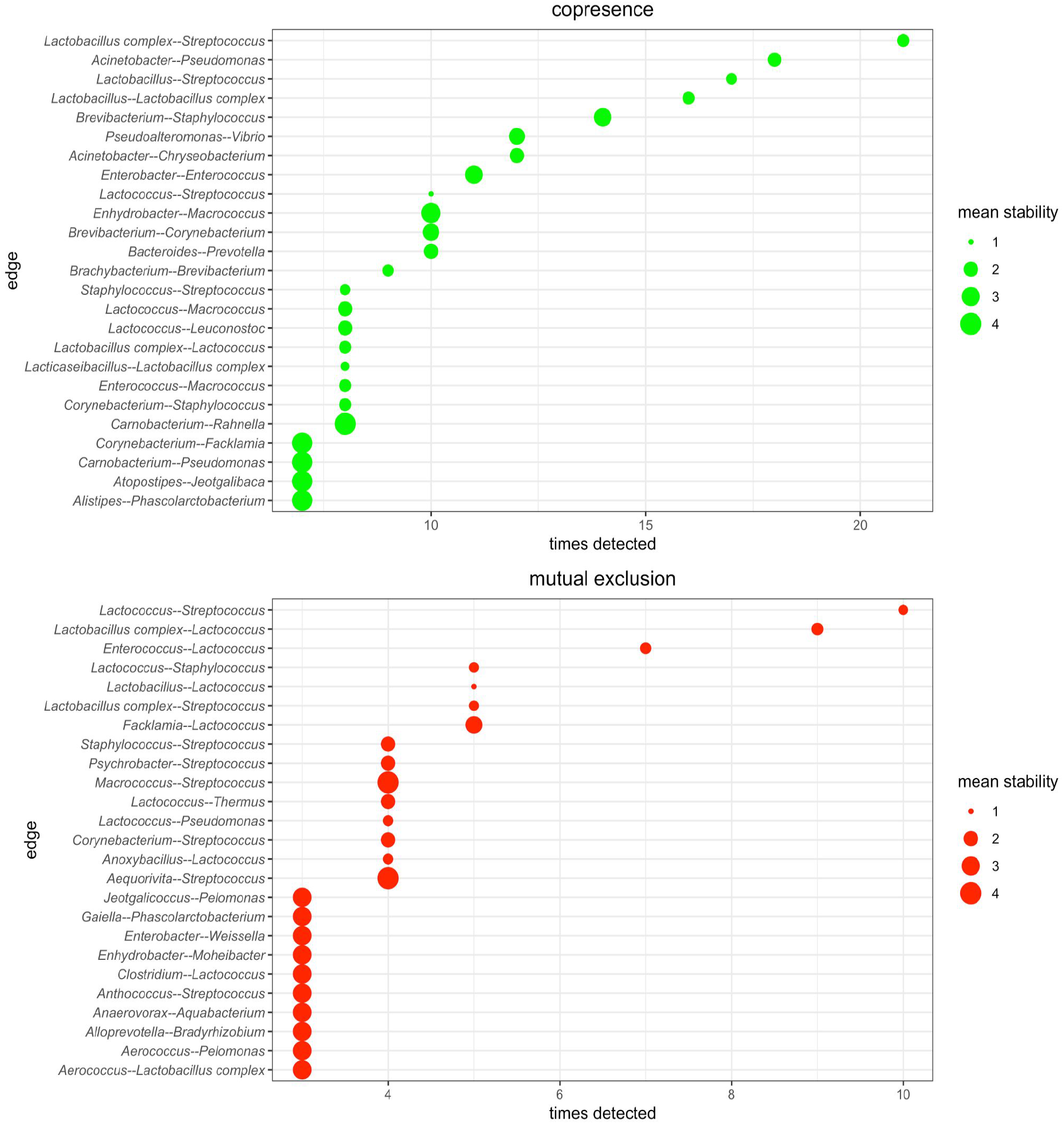
The 50 topmost (in terms of frequency of detection, the maximum absolute frequency of detection is 140, given the fact that there were 35 studies analyzed by 4 inference methods) copresence and mutual exclusion association in cheese studies extracted from DairyFMBN (Supplementary Table 1).

The results are in good agreement with our previous study (Parente et al., 2018) with copresence associations usually observed when both members were either beneficial (starter or non-starter) or non-beneficial (spoilage or contaminants), while mutual exclusion associations were frequently observed in couples including one beneficial and one non-beneficial microorganism. Several associations match well those reported in the literature (Blaya et al., 2017; Mayo et al., 2021). Some mutual exclusion associations observed at low frequency include host associated (*Alloprevotella*, *Anaerovorax, Bacteroides, Phascolarctobacterium, Prevotella*) or prevalently environmental (*Aquabacterium, Bradyrhizobium, Enhydrobacter, Gaiella, Jeotgalicoccus, Pelomonas*) as found by exploring the Florilege database (http://migale.jouy.inra.fr/Florilege/), and may simply reflect different environmental niches within one or few studies. A similar situation is observed when only associations inferred by method SPIEC-EASI are considered (Supplementary Figures 13 and 14). Unfortunately, due to the sparsity of SPIEC-EASI networks only very few copresence associations are detected in three or more studies. On the other hand, several copresence associations have relatively high values of the edge betweenness quantile (i.e. they rank relatively high for this edge centrality measure, which identifies edges whose removal might destruct the network and possibly impair the functionality of the microbial community.

Unfortunately, although inferring networks at the genus level is the only viable strategy when studies using target regions of different length have to be compared, it provides very little information on which species of the genera for which the associations are detected. In our combined dataset several studies targeted 16S RNA or RNA gene regions of sufficient length to allow identification at the species level for a significant number of sequences. Therefore, we inferred microbial association networks without taxonomic aggregation and selected only associations in which both members were identified at the species level. The 50 topmost associations in terms of frequency of detection are shown in Figure 6, while details of the study in which the association was detected are provided in Supplementary Table 2.

**Figure 6.**
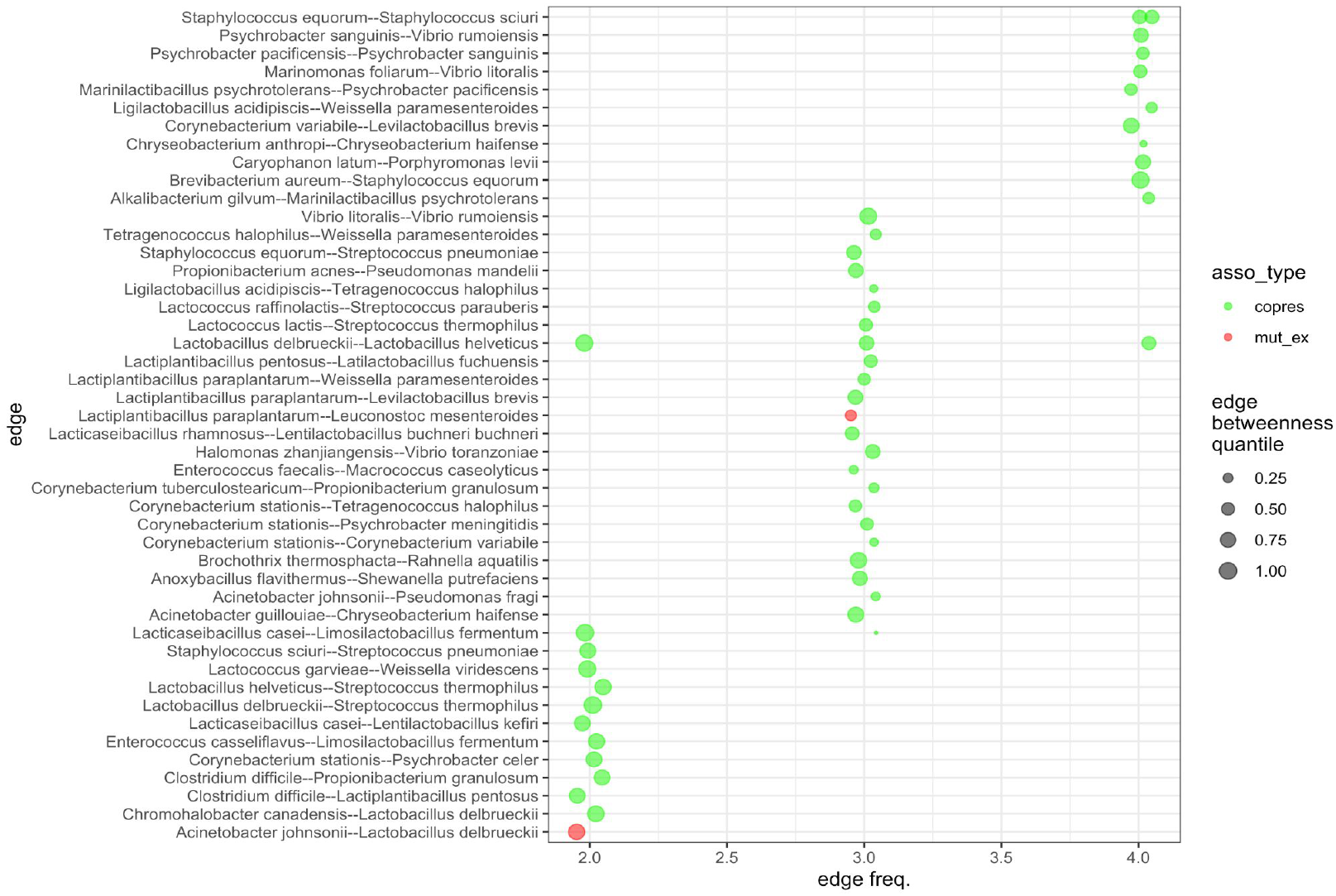
The 50 topmost copresence and mutual exclusion associations inferred by four different methods at the species level in cheese studies extracted from DairyFMBN (Supplementary Table 1). The number of methods detecting a given association is shown on the x axis, while a small random jitter was added to show when an association was detected in more than one study. The size of the points is made proportional to the edge betweenness quantile.

Given the relatively low number of associations inferred at the species level (compared to those inferred at the genus level), only three associations were detected in more than one study (*Staphylococcus equorum -- Staph. sciuri; Lactobacillus delbrueckii -- Lactob. helveticus; Lacticaseibacillus casei -- Limosilactobacillus fermentum*). However, the majority of the associations was detected by three inference methods or more in at least one study and might be therefore relatively robust. Most associations were detected in only a few studies (Bassi et al., 2015; Calasso et al., 2016; Cremonesi et al., 2020; De Filippis et al., 2016; Dolci et al., 2014; Dugat-Bony et al., 2016; Kamimura et al., 2019; Marino et al., 2019; Stellato et al., 2015) (see Supplementary Table 2). Only two mutual exclusion associations are included, although more have been detected by at least 2 methods. In general, the pattern observed at the genus level (copresence relationships between two beneficial or two non beneficial bacteria) is replicated. The interactions at the species level are much easier to test experimentally, to see if they merely reflect co-occurrence on the same microhabitat, or spatial or temporal niche or if they reflect true proto-cooperative or commensalistic relationships. Again, given the repeatability of our approach, as more data are added to DairyFMBN, more conserved associations at the species level could be added to this catalog. Indeed, it would be relatively easy to extend the analysis to other dairy related studies, including those on fermented milks or raw milk, and, in principle, a consensus network representing the dairy bacteria interactome could be constructed.

Finally, we decided to test the hypothesis of taxonomic assortativity (Kurtz et al., 2015) using a different approach compared to that used in a previous paper (Parente et al., 2018). The results for the data agglomerated at the genus level are shown in Supplementary figure 15. Even if the odds ratio for the occurrence of a copresence association among members of the same family was very high for the majority of studies, only in a few cases we were able to discard the null hypothesis that there was no difference in the probability of occurrence of a copresence association when the two nodes belonged to the same family or to different families.

## 4. A short note on phage-bacteria interaction networks

Bacteriophage infections are the main cause of failure of cheese fermentations whenever defined starter cultures are used (Erkus et al., 2013; Zotta et al., 2021) and have a dramatic impact on the structure and dynamics of bacterial populations and communities even when the more phage-tolerant undefined starter cultures are used (Erkus et al., 2013). In the latter case they have been found to be important in maintaining the equilibrium in complex association by a “kill the winner” mechanism (Erkus et al., 2013; Flores et al., 2011), which prevents the elimination from cultures reproduced by back-slopping of the slow-growing strains, by bacteriophage killing of the most competitive strains.

The statistical structure of Phage-Bacteria Interaction Networks (PBINs), which are typically represented as bipartite networks, has been elucidated (Flores et al., 2011; Weitz et al., 2013), and differences between the structure of PBINs for *Lactococcus* (which tend to show nestedness) and *S. thermophilus* (which tend to be modular) have been attributed to the differences in phage resistance mechanisms in these species.

Surprisingly enough, to the best of our knowledge, approaches based on bipartite network analysis have not been recently used to elucidate the structure of PBINs of dairy LAB. The development of new high-throughput approaches for the study of the metavirome in dairy products (see Zotta et al., 2021 for a recent review) combined with network science approaches is certainly promising in studying the evolution of bacteriophage-host relationships in natural starters and cheese in self-assembled communities (Canon et al., 2020) in cheese manufacture, like the many cheeses produced by using undefined starters reproduced by back-slopping (Zotta et al., 2021), in which dispersal, diversification, evolution and drift (Nemergut et al., 2013) all play a role in shaping the structure, dynamics and function of the cheese microbial communities, at least in early stages of cheese-making. Finally, beyond their role in regulating the structure of bacterial populations, bacteriophages (and prophages) of LAB may have other beneficial effects in microbial communities (induction of lysis, with release on nutrients for other members of the community; providing mechanisms for recombination and gene exchange; Paillet and Dugat-Bon, 2021).

Use of the concepts and computational approaches of bipartite network analysis (available, for example in the R bipartite package; Dormann et al., 2008) may be of assistance to both scientists interested in studying the structure and evolution of PBINs in cheese and to starter companies seeking to develop phage rotation schemes by facilitating the identification of potential hub strains and the identification of modules of virulent phages and susceptible strains.

## 5. Future prospects

### 5.1 Does inference of microbial association networks provide useful information?

Even with all limitations related to the inference of association networks from metataxonomic data (see section 2 and Röttjers and Faust, 2018 for a review), microbial association network inference offers invaluable information which may then be used to guide experiments to confirm the nature of the interactions confirmed in silico. The availability of simple to use workflows to calculate network properties and to formally compare networks (Peschel et al., 2020) can be used to formally test hypotheses on the effect of conditions (heat treatment, spoilage, use of starter cultures, etc.) on the structure of association networks. The networks structure of host microbiomes is known to be dramatically affected by disease (Layeghifard et al., 2017; Liu et al., 2020; Ma, 2018) and this has been proven to be true for mastitis (see Parente et al., 2020 for a review). Analysis of microbial association networks may help to mine for previously unknown interactions among bacteria relevant for cheese ripening and provide the basis for the design of microbiome based starter cultures (Canon et al., 2020; Mayo et al., 2021).

### 5.2 Down to the strain level

One limitation of HTS targeting 16S rRNA gene is that, at the very best, they can provide taxonomic resolution at the genus and, sometimes, at the species level. However, when Amplicon Sequence Variants are inferred or when improved taxonomic databases are used, taxonomic resolution can be improved (Pujato et al., 2018). On the other hand, biological interactions in cheese happen at the strain level, not at the species level (i.e. different strains of the same species can show a different interaction pattern). This is certainly true for bacteriophage-host interactions (Erkus et al., 2016, 2013) and has also been recently demonstrated for surface ripened cheeses (Niccum et al., 2020). As mentioned above (see 1.3 and 3), bacteriophages strongly affect the structure and microdiversity of mixed strain starters, through a “kill the winner” mechanism that prevents fast growing strains to take the dominance, especially when back-lopping is used for propagation. Strain specific behaviours and response to positive (indirect effects due to change in pH; direct effects due to cross-feeding relationships possibly related to the availability of iron, vitamins, amino acids) and negative (amensalism, competition) relationships in simplified communities including different combinations of different strains of *Staphylococcus equorum*, *Brachybacterium alimentarium, Brevibacterium aurantiacum* and a *Penicillium* strain), in simulated cheese-rind experiments, result in different community assemblies, which are reflected by changes in quality relevant feature (volatiles, color). Strain specific relationships were also found in interactions mediated by volatile compounds (Cosetta et al., 2020).

Although HTS methods targeting protein-coding genes have been used to study the structure of population of key bacterial species (see Bertuzzi et al., 2018; Moser et al., 2018; Walsh et al., 2020 for a recent review), they can only detect sequence variants of selected genes, not strains. Shotgun metagenomic approaches can provide full resolution for both bacterial strains and bacteriophages (Afshari et al., 2020, 2018; Walsh et al., 2020). However, the cost and the need for computing resources is still significantly higher compared to amplicon targeted approaches, and, as a result, shotgun metagenomic studies typically have a lower number of samples and the relatively low diversity of cheese microbial communities may prevent the detection of the less abundant members of the microbiota. This situation is likely to change in the future and integration between metagenomics, metatranscriptomics and metabolomics, and custom designed qPCR methods to quantitatively monitor single strains or lineages may offer great potential for the study of ecological and metabolic interaction in cheese (Erkus et al., 2013; Niccum et al., 2020).

### 5.3 Model based approaches and effect of environmental variables

Correlation or conditional dependence tools may be unable to detect some negative interactions (like amensalism; Weiss et al., 2016), especially in cross sectional studies based on compositional data, and may be unable to properly separate the effect of habitat filtering if environmental variables are not included. even if the number of time series studies using metataxonomic or meta-omic approaches for the study of the dynamics of cheese microbiota in increasing (see Supplementary Table 1 for some examples), the lack of quantitative measurements of the target genes and the lack of metadata related to significant environmental variables (temperature, pH, a_W_, salt in moisture) prevents the use of model based approaches for the inference of microbial interactions. In principle, using absolute quantification of the total amount of target (16S RNA or RNA gene) is possible, generally using qPCR or internal standards, and has been attempted in a few studies (Cauchie et al., 2017; Fougy et al., 2016; Rouger et al., 2018; Zotta et al., 2019), but only rarely attempts have been made to use these data to study the dynamics of members of the bacterial community (Zotta et al., 2019). In addition, when DNA is used as a target the question of the viability of the detected taxa remains open, and qPCR of 16S RNA or RNA gene can only provide partial indications of the real abundance of a given taxon, given the differences in copy numbers of these targets (Větrovský and Baldrian, 2013). On the other hand, lineage specific qPCR methods have successfully been used to study the dynamics of individual strains or groups of strains in cheese (Erkus et al., 2013, 2016). Therefore, in principle, use of time series metataxonomic data coupled to model based inference methods (such as MetaMis; Shaw et al., 2016) may be feasible and may be used to test in cheese model complex interactions after their preliminary evaluation in simplified laboratory models. More study is definitely needed in this area.

## Acknowledgements

This research did not receive any specific grant from funding agencies in the public, commercial, or not-for-profit sectors.

