## Supplementary Material for "Microbial association networks in cheese: a meta-analysis"

Scuola di Scienze Agrarie, Forestali ed Alimentari, Università degli Studi della Basilicata, Potenza, Italy

Supplementary material

#### Index

1. DATA AND SOFTWARE. 2
2. ANALYSIS WORKFLOW. 2
3. INFERENCE OF ASSOCIATION NETWORKS AT DIFFERENT TAXONOMIC LEVELS. 7
4. COMPARING INFERENCE METHODS. 8
5. COMPARING GLOBAL NETWORK PROPERTIES. 12
6. NODE PROPERTIES. 14
7. EDGE PROPERTIES. 14

REFERENCES 19

### 1. Data and software.

The metataxonomic data used for the inference of microbial association networks were extracted from DairyFMBN v2.1.6 (Parente et al., 2020, Parente, 2021), a specialized version of FoodMicrobionet (Parente et al., 2019), which is available on Mendeley Data (https://data.mendeley.com/datasets/3cwf729p34/5).
The list of studies used in the analysis is shown in Supplementary Table 1.

### 2. Analysis workflow.

A demo of the pipeline used for the inference and analysis of microbial association networks is available from GitHub (https://github.com/ep142/MAN_in_cheese) as a .Rmd notebook which, when run properly, will generate a ready to use report and a number of publication quality images and tables.
The analysis workflow is summarized in Supplementary Figure 1. Basically:

1. general options for plotting, saving and inference (including the inference methods to use) are set
2. an arbitrary number of phyloseq objects (McMurdie and Holmes, 2013) containing the data are imported in a list. This can be either objects extracted from DairyFMBN or phyloseq objects generated using a suitable bioinformatics pipeline. Study metadata are also imported in this stage as a tab delimited file;
3. samples with low number of sequences and studies with low number of samples are removed; calculation of diversity indices is also performed at this stage
4. taxonomic filtering (removal of Amplicon Sequence Variants identified as *Eukaryotes*, mitochondria or chloroplasts, removal of sequences identified above the family level) and agglomeration (no agglomeration, or agglomeration at the species, genus or family level) are performed using mostly functions from package phyloseq
5. prevalence and abundance filtering is performed to remove the least prevalent and abundant taxa (even if this can be performed within the netConstruct() function, see below); prevalence and abundance plots and tables are generated; reports on the effect of filtering on the number of taxa and sequences are generated;
6. network inference is performed for all datasets and, within dataset, all inference methods using the netConstruct() function of the NetCoMi package (Peschel et al., 2020); errors are trapped using try() and the results (as microNet or try objects) are put in a list and a report is generated
7. networks are analysed using netAnalyze() function; global network properties are extracted and joined with metadata and microbial diversity indices and evenness indices (including Pielou J and average Bray-Curtis dissimilarity, see Parente et al., 2018 for details)
8. node statistics (cluster membership and centrality indices) are then extracted from the microNetProps objects and taxonomic and prevalence and abundance information are merged
9. the networks are extracted ad tidygraph objects (Pedersen, 2020) and edge betweenness is calculated; Venn diagrams showing edges in common between different inference methods are optionally generated
10. global properties of the networks are compared using Principal Component Analysis
11. networks are plotted using ggraph (Pedersen, 2021) or NetCoMi and optionally compared in grids

12. node plots are generated for each dataset/inference method
13. stable edges (edges identified with more than one method within a given dataset or over different datasets) are identified
14. taxonomic assortativity is tested using epi.2by2() function of the epiR package (Stevenson et al., 2021)

Several aspects of the workflow can be personalized by setting options (locations of the input and output files, resolution of the graphs, filtering and taxonomic agglomeration options, network inference methods to be used and their parameters).

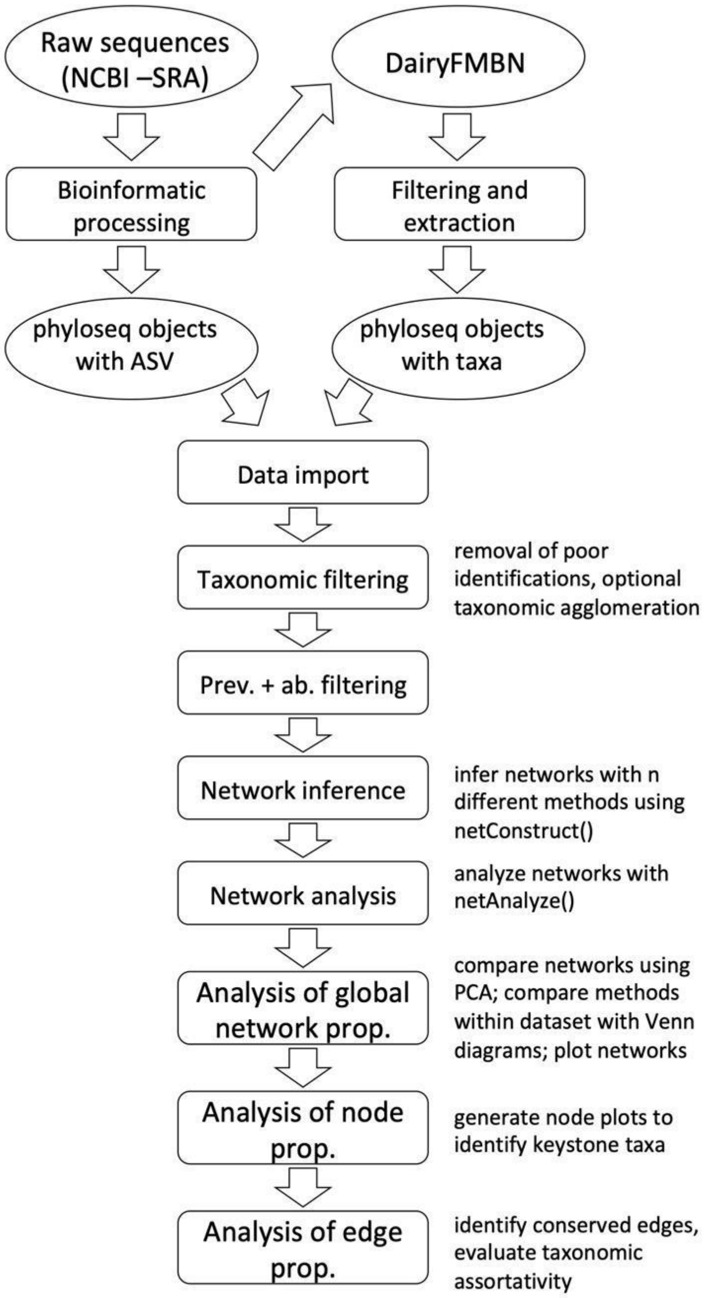


**Supplementary Figure 1**. Schematic representation of the workflow used in the meta-analysis.

#### Supplementary Table 1.

Details of the studies used for inference of microbial association networks. The studies were extracted from DairyFMBN 2.1.6 (Parente et al., 2020, Parente, 2021).

| Study | T arget | R egion | R ead length (bp) | NCBI SRA seq. accn. | S amples | Description | Type[1] | Design | Reference |
|---|---|---|---|---|---|---|---|---|---|
| ST1 | 16S RNA gene | V1-V3 | 504 | SRP052240 | 29 | Commercial high-moisture Mozzarella cheese produced with different acidification methods | cs (1) | descriptive | Guidone et al., 2016 |
| ST2 | 16S RNA gene | V1-V3 | 498 | SRP057506 | 24 | Undefined strain starters (milk cultures) for high-moisture Mozzarella cheese | cs (1) | descriptive | Parente et al., 2016 |
| ST3 | 16S RNA gene | V1-V3 | 465 | SRP033419 | 50 | Undefined strain starters (whey cultures) and cheese curds for water-buffalo Mozzarella, Grana Padano (and Parmigiano Reggiano cheese | cs (1) | 3x2 groups (small) | De Filippis et al., 2014 |
| ST6 | 16S RNA | V1-V3 | 485 | SRP038100 | 22 | Milk, curd and ewe's milk Canestrato cheese during ripening | ts (11) | time series + descriptive | De Pasquale et al., 2014a |
| ST8 | 16S RNA | V1-V3 | 490 | SRP040575 | 27 | Milk (from different lactation stages), curd and Fontina cheese from three different dairies | mx (3) | 3x2 groups (small) | Dolci et al., 2014 |
| ST9 | 16S RNA | V1-V3 | 469 | SRP044294 | 39 | Piedmont hard cheese made from raw milk: milk, curd and cheese throughout ripening | mx (7) | time series + descriptive | Alessandria et al., 2016 |
| ST10 | 16S RNA gene and 16S RNA | V1-V3 | 601 | SRP061555 | 67 | Caciocavallo Silano cheese manufacture: starter culture, milk, curd and cheese throughout ripening | ts (5) | 2x2x2 groups (small) | De Filippis et al., 2016 |
| ST18 | 16S RNA gene | V1-V3 | 454 | SRP058584 | 45 | Environmental swabs from an Italian dairy plant and different kind of cheeses (Mozzarella; Ricotta; Scamorza; Caciocavallo; Grancacio) produced in the same plant | cs (1) | time series + descriptive | Stellato et al., 2015 |
| ST22 | 16S RNA gene | V1-V3 | 465 | -- | 48 | Environmental swabs from an Italian dairy plant and different kind of cheeses (Caciotta; Caciocavallo Pugliese) and cow milk produced in the same plant | mx (10) | 2 groups | Calasso et al., 2016 |
| ST23 | 16S RNA gene | V3-V4 | 398 | SRP055798 | 37 | Grana Padano cheese samples with or without blowing defect, from different factories, at different ripening | cs (1) | descriptive | Bassi et al., 2015 |

| | | | | | | | | | |
|---|---|---|---|---|---|---|---|---|---|
| | | | | | | times, produced with and without lysozyme | | | |
| ST25 | 16S RNA gene | V4-V5 | 232 | ERP009223 | 31 | Continental (Swiss type cheese) produced early and late in the day, core and rind samples included, different ripening times | cs (1) | descriptive | O'Sullivan et al., 2015 |
| ST32 | 16S RNA gene | V4 | 500 | -- | 93 | Artisanal soft, semi-hard and hard cheeses from raw or pasteurized cow, goat, or sheep milk | cs (1) | descriptive, possibly 3 groups | Quigley et al., 2012 |
| ST33 | 16S RNA gene | V4 | 500 | ERP006630 | 58 | Investigating the role of the microbiota in Pink Cheese. Cheddar, Emmental and cheese coloured with Annatto, either unspoiled or spoiled. | cs (1) | 2x2 groups small | Quigley et al., 2016 |
| ST34 | 16S RNA | V3-V4 | 422 | SRP060430 | 46 | Bovine ricotta cheese (two lots, winter and spring) without or with pink discoloration, throughout storage at 8°C. | mx (7) | 2 groups small | Sattin et al., 2016 |
| ST36 | 16S RNA | V1-V3 | 424 | SRP110830 | 50 | Raw cow milk and burrata cheese protective lactobacilli cheese with or without dietary fibres an | mx (4) | 3 groups, small | Minervini et al., 2017 |
| ST39 | 16S RNA gene | V3-V4 | 435 | SRP126475 | 48 | Teat skin, raw cow milk and Cantal cheese) microbiota, as a function of grazing system | mx (3) | 4+ groups, small | Frétin et al., 2018 |
| ST41 | 16S RNA gene | V3-V4 | 425 | SRP071345 | 95 | Microbiota of core and rind of 12 French cheeses | cs (1) | 4+ or 2 groups, small | Dugat-Bony et al., 2016 |
| ST43 | 16S RNA gene | V1-V3 | 420 | SRP051167 | 26 | Cheese with brown defect and cheese environment | cs (1) | 4 groups (small) | Guzzon et al., 2017 |
| ST44 | 16S RNA gene | V1-V3 | 371 | SRP070077 | 38 | Caciocavallo cheese, throughout ripening, with milk obtained under different cow's feeding regimes | mx (7) | 2 groups (small) | Giello et al., 2017 |
| ST45 | 16S RNA gene | V3-V4 | 463 | SRP103624 | 92 | Gouda cheese (15 brands), from pasteurized or raw milk. Spatial (core, surface, middle) variability was assessed. Cheese age (2-18 mo.) confounded with brand. | cs (1) | 2 groups | Salazar et al., 2018 |
| ST48 | 16S RNA gene | V3-V4 | 427 | SRP156292 | 39 | High moisture Mozzarella cheese (cow or buffalo milk), produced with different types of starters | cs (1) | descriptive | Marino et al., 2019 |
| ST49 | 16S RNA gene | V3-V4 | 412 | SRP165151 | 196 | Artisanal raw milk cheeses from Brazil (11 different types from 5 geographical areas) | cs (1) | descriptive | Kamimura et al., 2019 |
| ST74 | 16S RNA gene | V3 | 151 | SRP170819 | 375 | Cow milk, cheese (4 types: Brie, Cheddar, Gruyere, Jarlsberg) and environmental samples, from farm to fork for an artisanal cheese production facility | cs (1) | descriptive | Falardeau et al., 2019 |
| ST106 | 16S RNA | V1-V3 | 369 | SRP059382 | 30 | Active microbiota from Italian PDO ewe's cheese (Pecorino toscano; Pecorino siciliano; Fiore sardo) | cs (1) | descriptive | De Pasquale et al., 2016 |
| ST110 | 16S RNA gene | V3-V4 | 427 | SRP212264 | 112 | Dynamics of the microbiota of semisoft caciotta cheese | mx (3) | 4+ | Calasso et |

| | | | | | | | | | |
|---|---|---|---|---|---|---|---|---|---|
| | | | | | | produced a washed rind protocol, with or without attenuated adjuncts and surface inoculants | | | al., 2020 |
| ST115 | 16S RNA gene | V4 | 245 | SRP233045 | 63 | Microbiota of core and rind of Cheddar, Provolone and Swiss type cheese produced in Oregon from pasteurized milk | cs (1) | 3x2 groups, small | Choi et al., 2020b |
| ST131 | 16S RNA gene | V4 | 245 | SRP244702 | 108 | Microbiota of Cheddar cheese during production and aging (26 months). Two batches, raw and pasteurized milk included | ts (24+) | -- | Choi et al., 2020a |
| ST136 | 16S RNA gene | V3-V4 | 427 | SRP256471 | 47 | Microbiota of F romadzo cheese (a semi-hard, ripened cheese from Valle d'Aosta) made from milk from two different farms during ripening | mx (4) | 2x2 groups | Dolci et al., 2020 |
| ST146 | 16S RNA gene | V3-V4 | 427 | ERP119763 | 22 | Microbiota of Paipa cheese, a Colombian semi-ripened cheese made from raw cow milk | mx (4) | descriptive | Castellanos -Rozo et al., 2020 |
| ST148 | 16S RNA gene | V3-V4 | 427 | SRP274316 | 67 | Bacterial microbiota of Parmigiano Reggiano throughout cheese ripening (curd-24 mo.) for 6 production batches | mx (5) | descriptive | Bottari et al., 2020 |
| ST149 | 16S RNA gene | V3-V4 | 425 | SRP254987 | 102 | Microbiota of raw cow bulk milk and vat milk, natural whey culture and Trentingrana as affected by chlorine wash of milking equipment | mx (3) | possibly 3 groups | Cremonesi et al., 2020 |
| ST150 | 16S RNA gene | V3-V4 | 426 | SRP257637 | 58 | Evolution of microbiota during production of Robiola (milk, natural milk starter, cheese) di Roccaverano, an artisanal Protected Designation of Origin soft cheese made with raw goat milk by addition of a natural milk starter (NMS), from the Piedmont region of Italy | mx (3) | possibly 4 groups | Biolcati et al., 2020 |
| ST157 | 16S RNA gene | V4 | 271 | SRP290895 | 40 | Bacterial microbiota of commercial Australian Cheddar cheese, three brands, at different ripening ages | cs (1) | possibly 3 groups | Afshari et al., 2020 |
| ST165 | 16S RNA gene | V3-V4 | 425 | ERP121277 | 45 | Bacterial microbiota of Pélardon cheese, a French PDO goat milk cheese with white bloomy rind, during ripening | mx (3-7) | | Penland et al., 2021 |

1 cs cross sectional study; ts time series (one or very few cheese makings followed over a relatively long period of time); mx mixed (several cheeses with some time points available for each). The number of time points is shown in parentheses.

#### 3. Inference of association networks at different taxonomic levels.

Microbial association networks were inferred for two studies, ST49 and ST136 (Supplementary Table 1) using the phyloseq objects with Amplicon Sequence Variants (no taxonomic agglomeration, prior to merging in FoodMicrobionet, Parente et al., 2019), with taxonomic agglomeration at the lowest taxonomic level (this is the level of taxonomic agglomeration of objects stored in DairyFMBN) and after taxonomic agglomeration at the genus level. Prevalence and abundance filtering were performed by removing ASVs or "taxa" which had a prevalence <0.05 and a relative abundance <0.005. Inference of networks was performed using CCREPE and SPIEC-EASI with neighbourhood selection, the networks were transformed in tidygraph objects and plotted using ggraph. The networks with taxonomic aggregation at the lowest possible level are shown in Figure 2 of the paper, while those at the ASV and genus level are shown in Supplementary Figures 2 and 3 respectively.

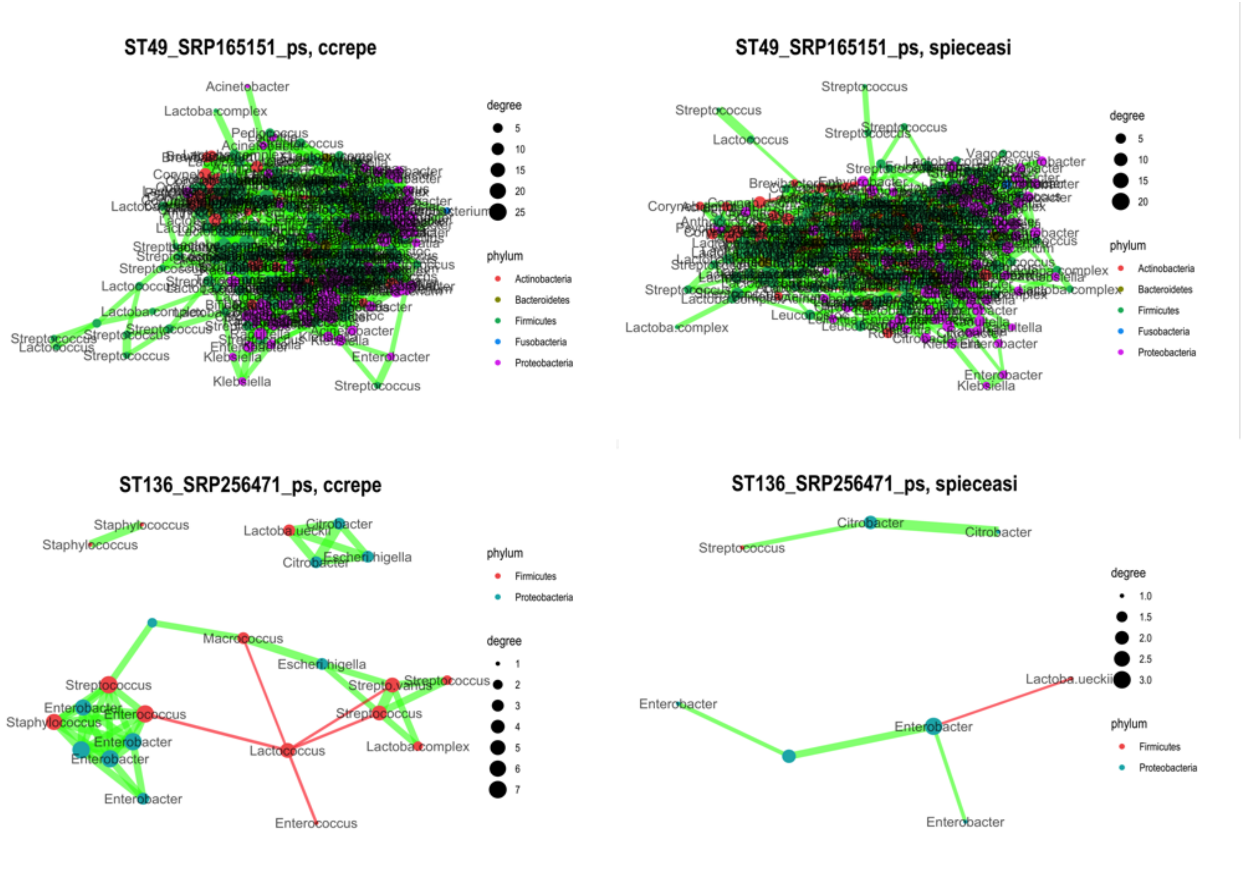


**Supplementary Figure 2.** Microbial association networks inferred Amplicon Sequence variants for studies ST49 (Kamimura et al., 2019) and ST136 (Dolci et al., 2020) using methods CCREPE (Faust et al., 2012) or SPIEC-EASI (Kurtz et al., 2015). For each pane, the colour of nodes is determined by phylum, and the size by degree; the colour of the edges is red for mutual exclusion relationships and green for presence relationships; the thickness of the nodes is determined by the strength of the association measure, as determined in NetCoMi (Peschel et al., 2020). A force-based layout (Fruchterman–Reingold) was used for positioning nodes and edges. The name of the nodes has been abbreviated to 15 characters.

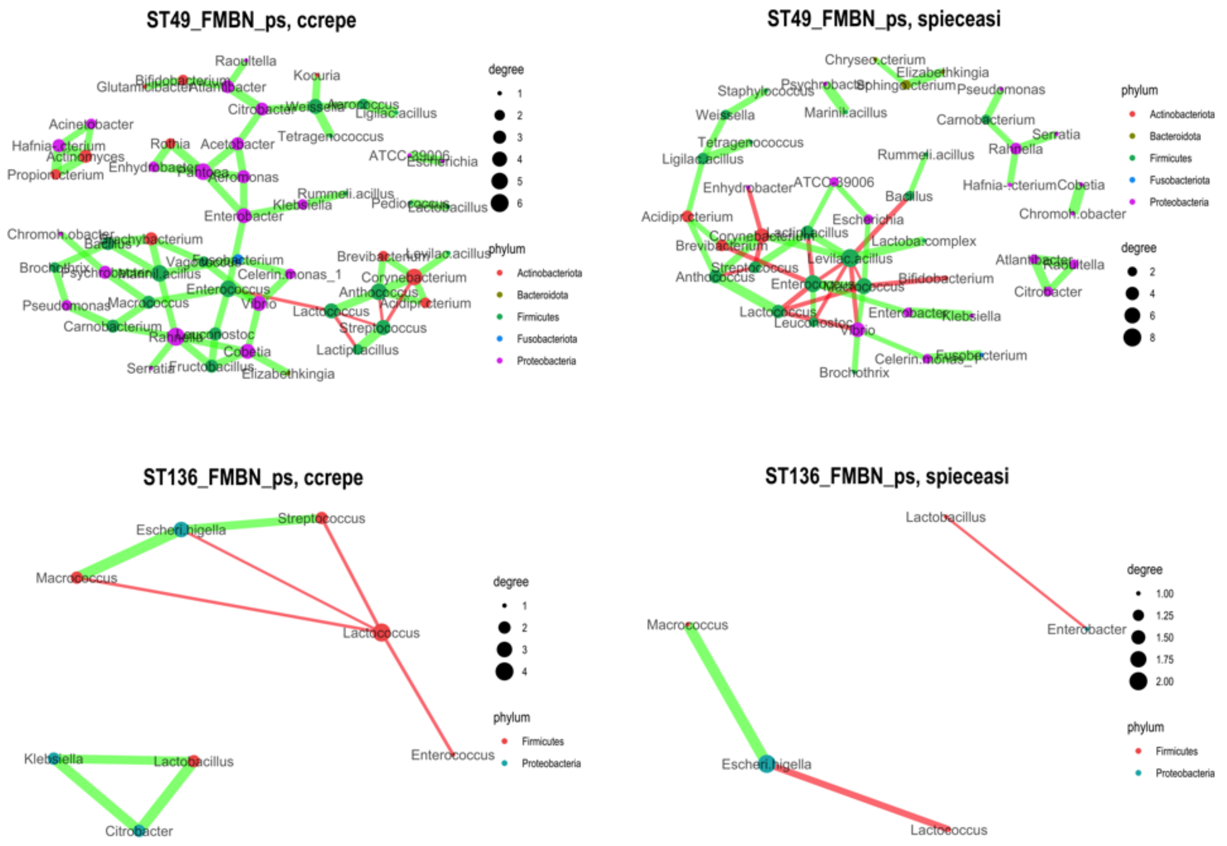


**Supplementary Figure 3.** Microbial association networks inferred from studies ST49 (Kamimura et al., 2019) and ST136 (Dolci et al., 2020) extracted from DairyFMBN 2.1.6 (https://data.mendeley.com/datasets/3cwf729p34/5), with taxonomic agglomeration at the genus level using methods CCREPE (Faust et al., 2012) or SPIEC-EASI (Kurtz et al., 2015). For each pane, the colour of nodes is determined by phylum, and the size by degree; the colour of the edges is red for mutual exclusion relationships and green for presence relationships; the thickness of the nodes is determined by the strength of the association measure, as determined in NetCoMi (Peschel et al., 2020). A force-based layout (Fruchterman–Reingold) was used for positioning nodes and edges. The name of the nodes has been abbreviated to 15 characters.

#### 4. Comparing inference methods.

Microbial association networks were inferred using four methods (SparCC, Sparse Correlations for Compositional data, Friedman and Alm, 2012; CCREPE Compositionality Corrected by REnormalization and PErmutation, Faust et al., 2012; SPIEC-EASI, SParse InversE Covariance Estimation for Ecological Association Inference, Kurtz et al., 2015; SPRING, SemiParametric Rank-based approach for INference in Graphical model, Yoon et al., 2019) on 35 datasets from 34 studies from DairyFMBN 2.1.6 (Supplementary Table 1) using a R workflow based mostly on R packages phyloseq, NetCoMi, and tidygraph (see section 2). Taxonomic agglomeration was performed at the genus level using the function tax_glom() of package phyloseq.

The inferred networks for two studies (ST41, Dugat-Bony et al., 2016; ST131 Choi et al., 2020a; see Supplementary Table 1 for details) are shown in Supplementary Figures 4 and 5.

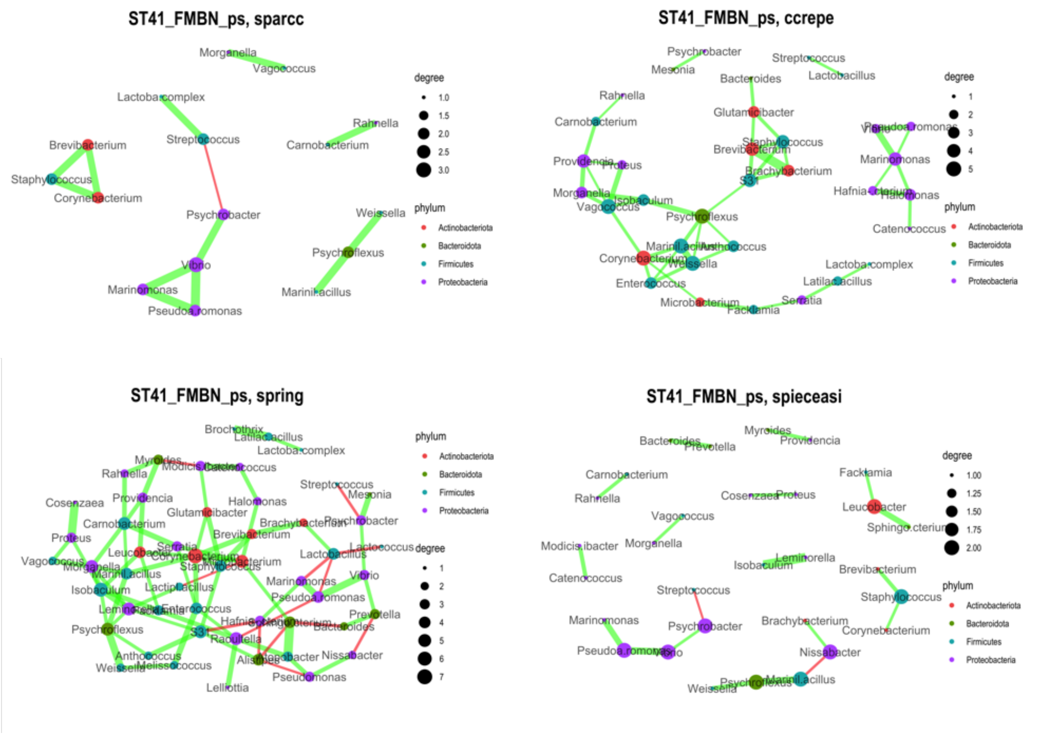


**Supplementary Figure 4**. Microbial association networks inferred at the genus level for studies ST31 (Dugat-Bony et al., 2019) using four methods (SparCC, Sparse Correlations for Compositional data, Friedman and Alm, 2012; CCREPE Compositionality Corrected by REnormalization and PErmutation, Faust et al., 2012; SPIEC-EASI, SParse InversE Covariance Estimation for Ecological Association Inference, Kurtz et al., 2015; SPRING, SemiParametric Rank-based approach for INference in Graphical model, Yoon et al., 2019). For each pane, the colour of nodes is determined by phylum, and the size by degree; the colour of the edges is red for mutual exclusion relationships and green for presence relationships; the thickness of the nodes is determined by the strength of the association measure, as determined in NetCoMi (Peschel et al., 2020). A force-based layout (Fruchterman–Reingold) was used for positioning nodes and edges. The name of the nodes has been abbreviated to 15 characters.

A number of comparisons (number of edges, PEP - Positive Edge Proportion, density, clustering coefficient, modularity, all estimated using the netAnalyze() function of NetCoMi) between methods were carried out using the 35 datasets (2 separate datasets, one for DNA and one for RNA were available for study 10) listed in Supplementary Table 1. The significance of the differences was tested, when appropriate, using a pairwise Wilcoxon test.
The distribution of the number of edges for each method is compared in Supplementary Figure 6.
The distribution of the network density of the networks for each method is compared in Supplementary Figure 7.
The distribution of positive edge proportion of the networks for each method is compared in Supplementary Figure 8.
The distribution of global clustering coefficient and modularity of the networks for each method is compared in Supplementary Figure 9 and 10, respectively.

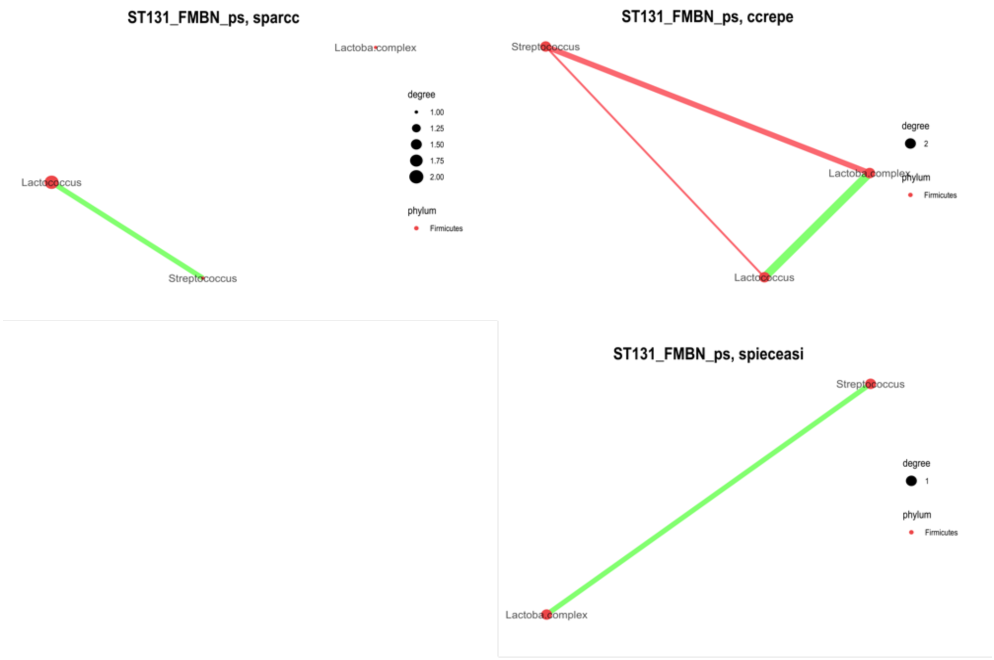


**Supplementary Figure 5.** Microbial association networks inferred at the genus level for studies ST141 (Choi et al., 2020a) using four methods (SparCC, Sparse Correlations for Compositional data, Friedman and Alm, 2012; CCREPE Compositionality Corrected by REnormalization and PErmutation, Faust et al., 2012; SPIEC-EASI, SParse InversE Covariance Estimation for Ecological Association Inference, Kurtz et al., 2015; SPRING, SemiParametric Rank-based approach for INference in Graphical model, Yoon et al., 2019). Only three methods returned networks with at least one edge. For each pane, the colour of nodes is determined by phylum, and the size by degree; the colour of the edges is red for mutual exclusion relationships and green for presence relationships; the thickness of the nodes is determined by the strength of the association measure, as determined in NetCoMi (Peschel et al., 2020). A force-based layout (Fruchterman–Reingold) was used for positioning nodes and edges. The name of the nodes has been abbreviated to 15 characters.

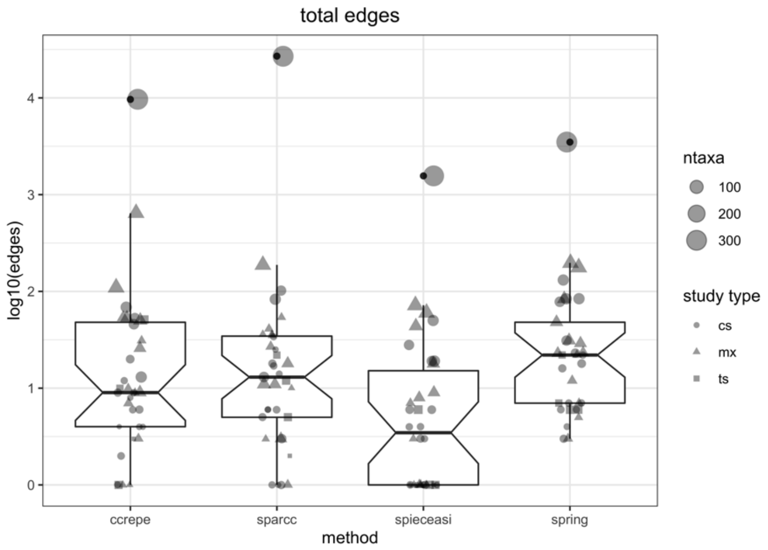


**Supplementary Figure 6.** A comparison of total number of edges in microbial association networks inferred after taxonomic agglomeration at the genus level for the studies listed in Supplementary Table 1.

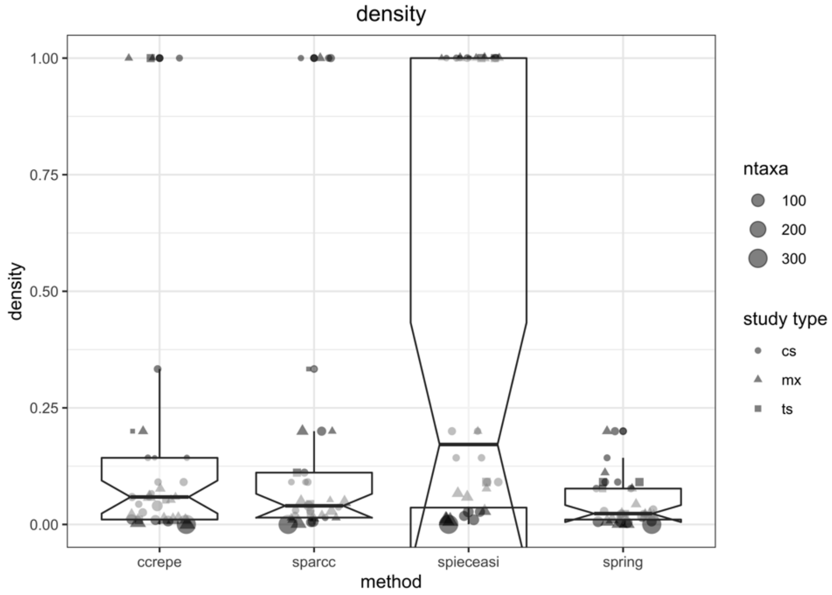


**Supplementary Figure 7.** A comparison of network density in microbial association networks inferred after taxonomic agglomeration at the genus level for the studies listed in Supplementary Table 1.

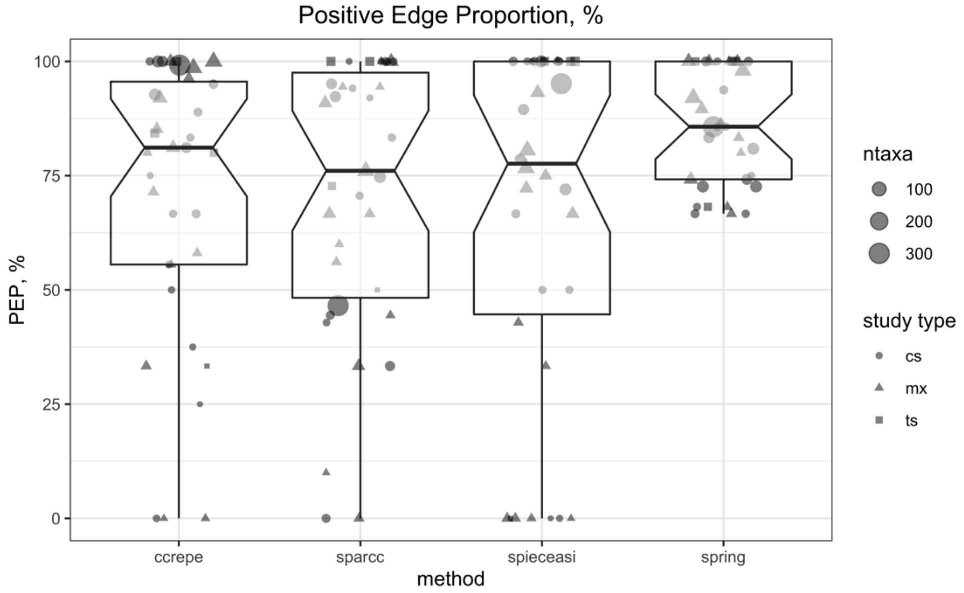


**Supplementary Figure 8.** A comparison of positive edge proportion in microbial association networks inferred after taxonomic agglomeration at the genus level for the studies listed in Supplementary Table 1.

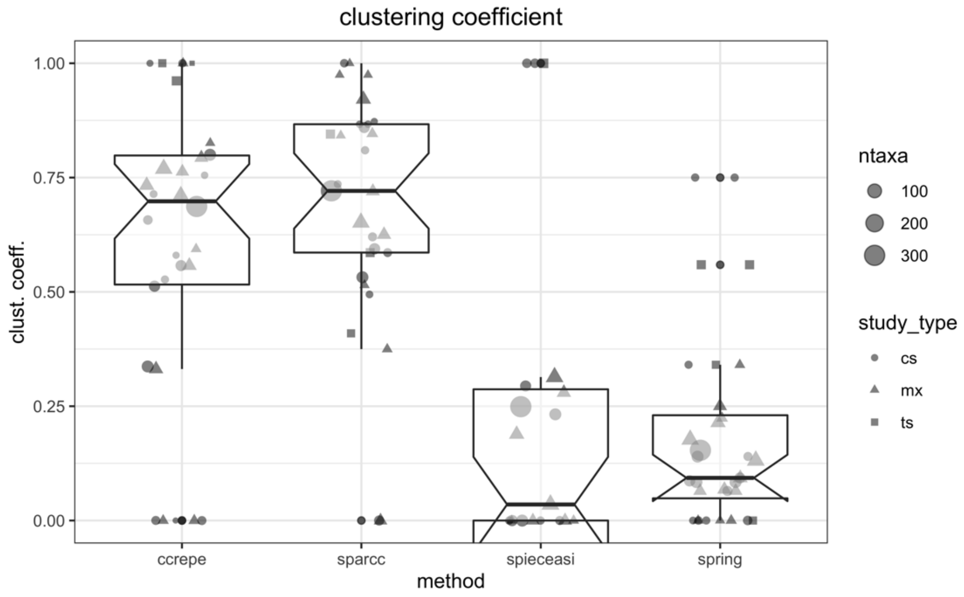


**Supplementary Figure 9.** A comparison of global clustering coefficient in microbial association networks inferred after taxonomic agglomeration at the genus level for the studies listed in Supplementary Table 1.

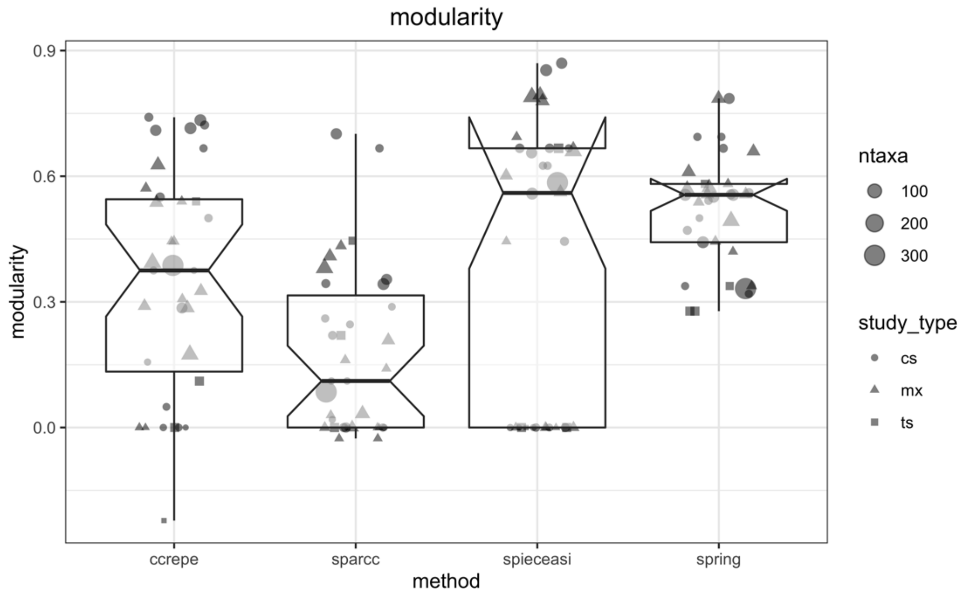


**Supplementary Figure 10.** A comparison of modularity in microbial association networks inferred after taxonomic agglomeration at the genus level for the studies listed in Supplementary Table 1.

#### 5. Comparing global network properties.

Microbial associations were inferred for all studies in Supplementary Table 1 after filtering agglomeration at the genus level using method SPIEC-EASI, and network properties were estimated using netAnalyze() function of NetCoMi (see Supplementary Figure 1) with the centrLCC parameter set to true. This resulted in calculation of centralities only for the largest connected component (LCC), i.e. when more than a connected component of the network was present (a group of nodes connected among them but not with other group of nodes in the same network), the centralities were only calculated for the LCC. Furthermore, the global network statistics were merged with metadata (number of taxa, types of study, diversity indices calculated as described in Parente et al., 2018 (Chao1, average Bray-Curtis distance, and an evenness index, Pielou J).

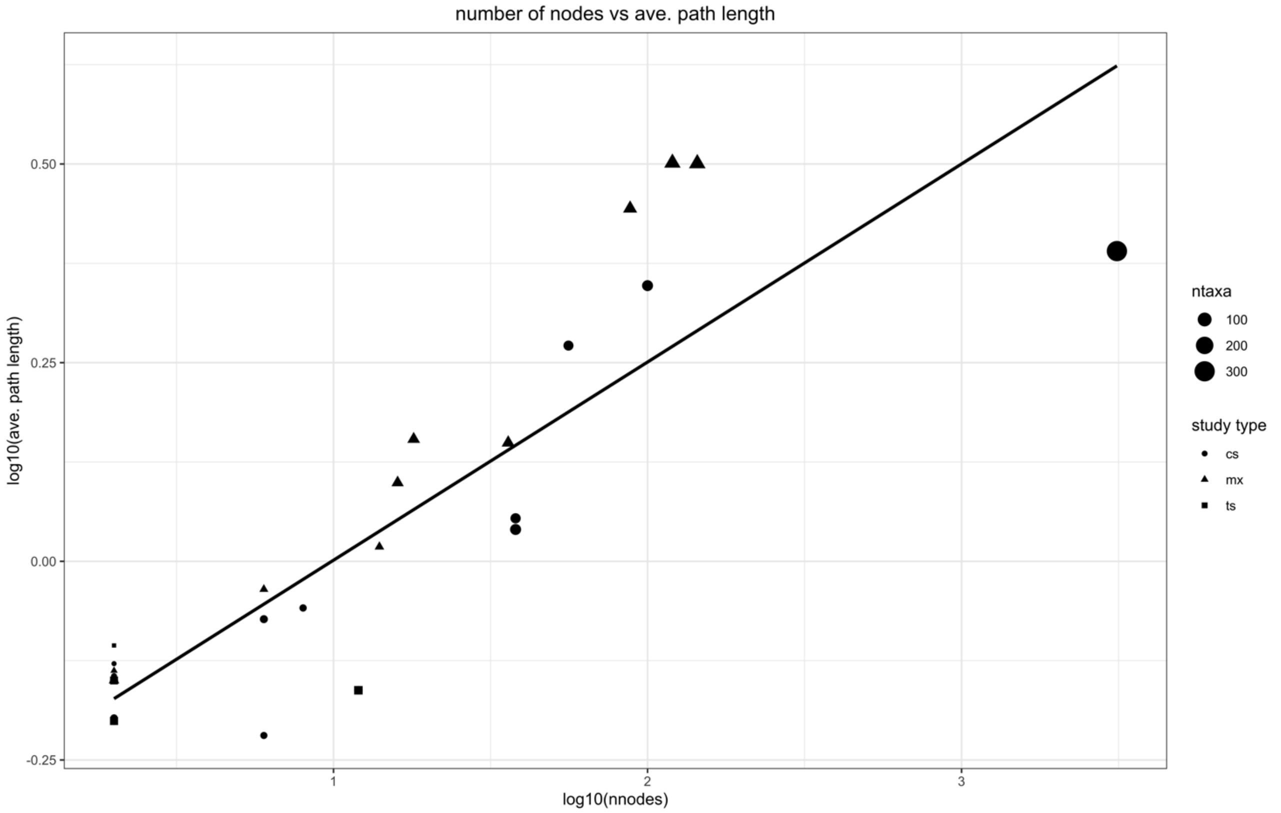


**Supplementary Figure 11.** Relationship between the size of the networks and the average path length for the largest connected component for microbial association networks inferred with method SPIEC-EASI after taxonomic agglomeration at the genus level for the studies listed in Supplementary Table 1.

The relationship between the number of nodes of the networks and the average path length of the largest connected component is shown in Supplementary Figure 11.
The correlation between selected variables, including network properties (average path length, avPath; density, calculated on all connected nodes, density_2; modularity; natural connectivity, i.e. the average eigenvalue of the adjacency matrix, a measure of the robustness of a graph, natConnect; number of connected nodes, nnodes; positive edge proportion, pep) and properties of the datasets (average dissimilarity avDiss: average of the Bay-Curtis distance matrix; average Chao1 index; Pielou J evenness) was calculated using the Pearson product moment coefficient. A principal component analysis with varimax rotation was carried out using function prcomp() and a biplot of the first two components was plotted using the autoplot() function of package ggfortify (Tang et al., 2016). Parallel analysis carried out using package psych (Revelle, 2020) confirmed that three components were sufficient to explain the variance (74%).

#### 6. Node properties.

Using the data generated in the previous session, node properties were extracted from the netProps objects created by netAnalyze() and further node properties (positive and negative degree, PEP) were calculated for each node. The results for all networks were combined into a data frame, merged with taxonomic metadata and used to generate graphs. An example for study ST49 (Kamimura et al., 2019) is shown in Supplementary Figure 12.

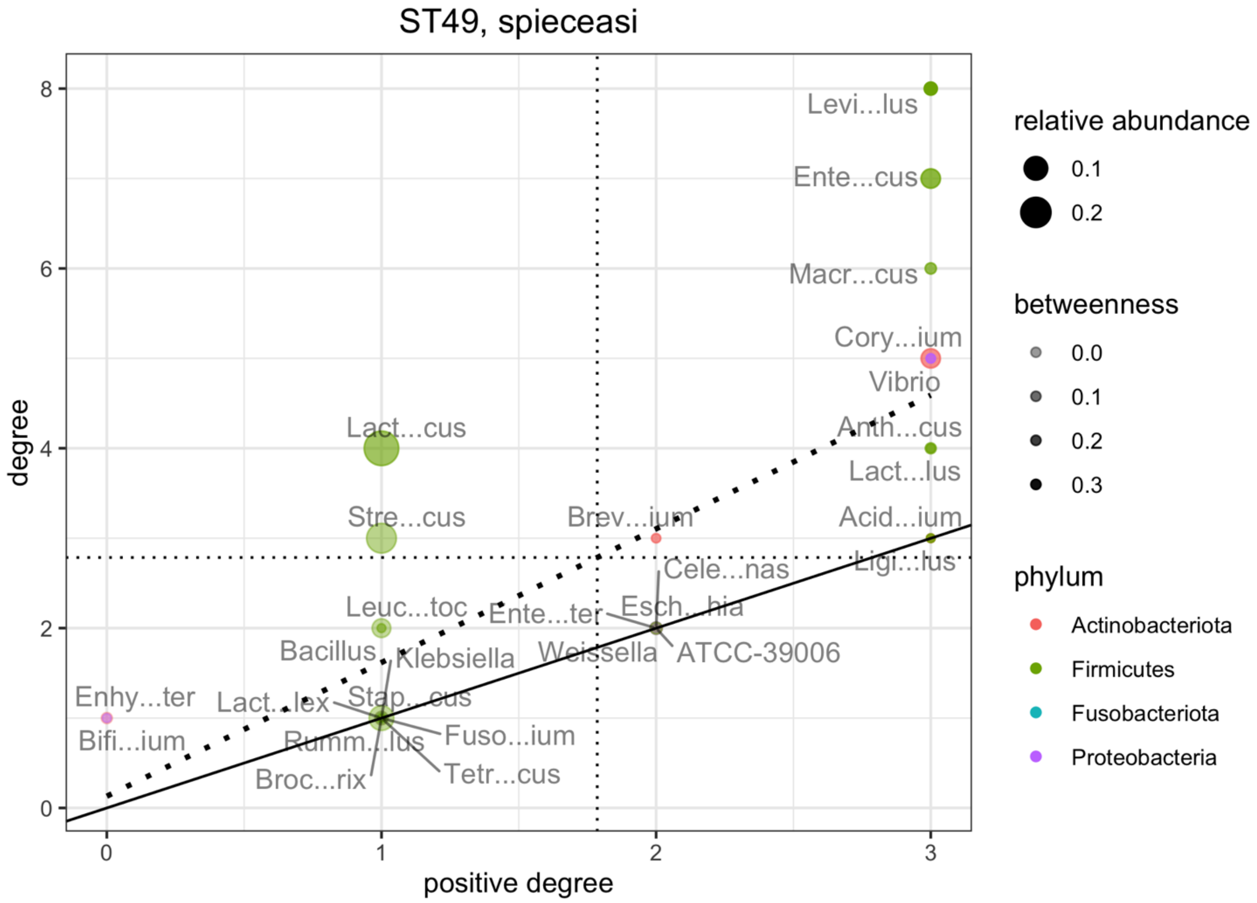


**Supplementary Figure 12**. Node plot for the microbial association network inferred using SPIEC-EASI after taxonomic agglomeration at the genus level for study ST49 (Kamimura et al., 2019). The name of the genera was abbreviated to 7 characters. The size of the nodes is made proportional to the relative abundance of the genus, and the transparency is made proportional to edge betweenness. The solid black line corresponds to equal values of degree and positive degree (i.e. all the nodes above have at least one negative edge). The dotted line is the linear regression line. The nodes above have a higher than average ratio of degree to positive degree.

#### 7. Edge properties.

Using the microbial association networks generated in section 4, we created a combined data frame with edges obtained from all methods. Taxonomic information was merged for both from and to nodes and flags showing if both nodes belonged to the same higher taxon (family, order or class) were created.

To evaluate which were the most stable and frequently detected nodes, the edges were first grouped by edge name and type (presence or mutual exclusion), pooling all data for different methods. The number of times and the frequency of detection of each node was calculated, together with the average frequency of detection within study (a measure of edge stability). The topmost 25 presence and mutual exclusion edges were plotted.
A similar procedure was used to evaluate which were the most frequently detected associations using method SPIEC-EASI. In this case summary statistics were calculated after grouping by edge. The topmost (in terms of absolute frequency of detection) 25 copresence and mutual exclusion associations are shown in Supplementary Figures 13 and 14, respectively.

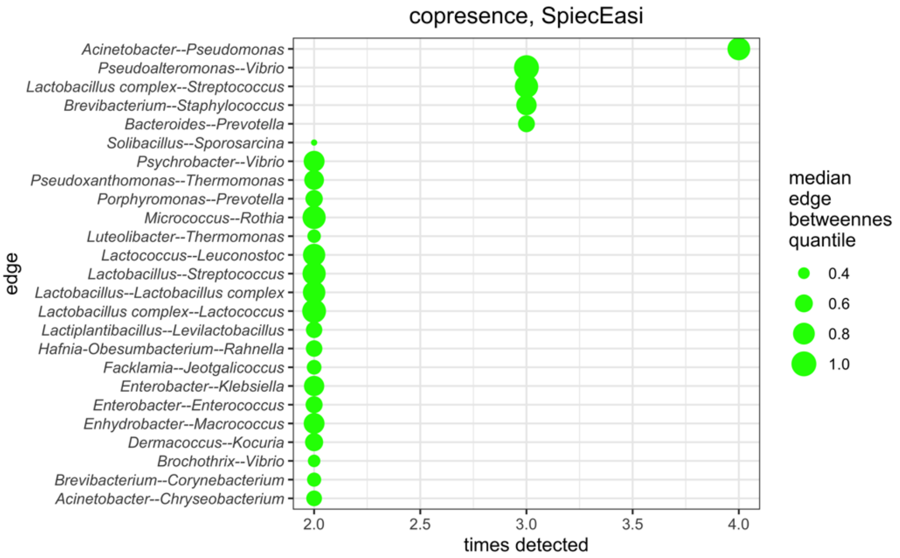


**Supplementary Figure 13.** The 25 topmost (in terms of number of studies in which the association was detected) copresence associations detected in microbial association networks inferred after aggregation at the genus level with method SPIEC-EASI. The size of the points is made proportional to the median edge betweenness quantile.

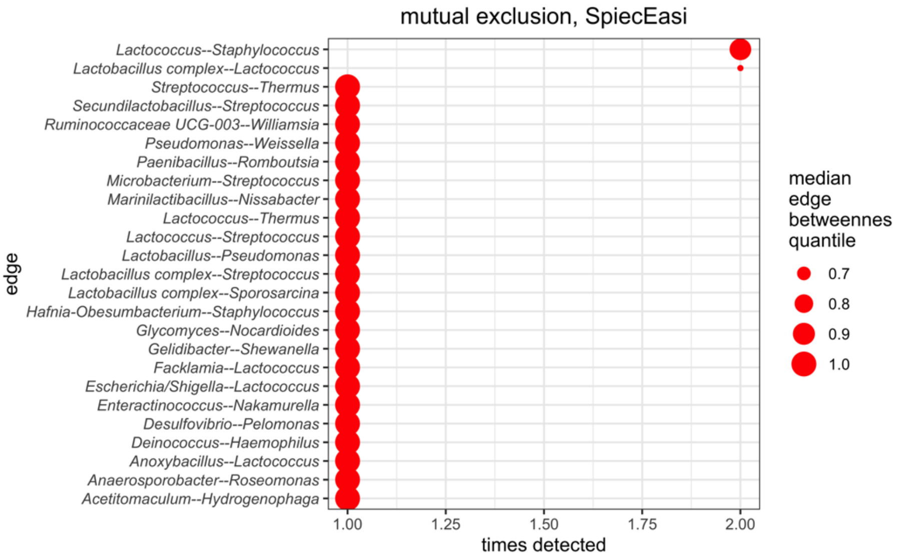


**Supplementary Figure 14.** The 25 topmost (in terms of number of studies in which the association was detected) mutual exclusion associations detected in microbial association networks inferred after aggregation at the genus level with method SPIEC-EASI. The size of the points is made proportional to the median edge betweenness quantile.

Finally, in order to obtain a preliminary evaluation of the most frequently detected associations at the species level, microbial association network detection was performed on the same set of studies with the four methods, without taxonomic agglomeration. Only edges in which both nodes had been identified at the species level were retained and the 50 most frequent associations were retained and plotted. The list of these edges is shown in Supplementary Table 2.
To evaluate if there was evidence of taxonomic assortativity, odds ratio (OR) and relative risk (RR) of having a co-presence association between members of the same higher taxon were calculated, after tabulation, using function epi.2by2() of package epiR (Stevenson et al., 2021). The decimal logarithm of odd ratio was calculated (using a dummy value of 5 whenever the OR was Inf) and a Yates test was testing the null hypothesis that being members of the same higher taxonomic group did not have any effect on the OR of having a presence relationship. The results for assortativity at the family level are shown in Supplementary Figure 15.

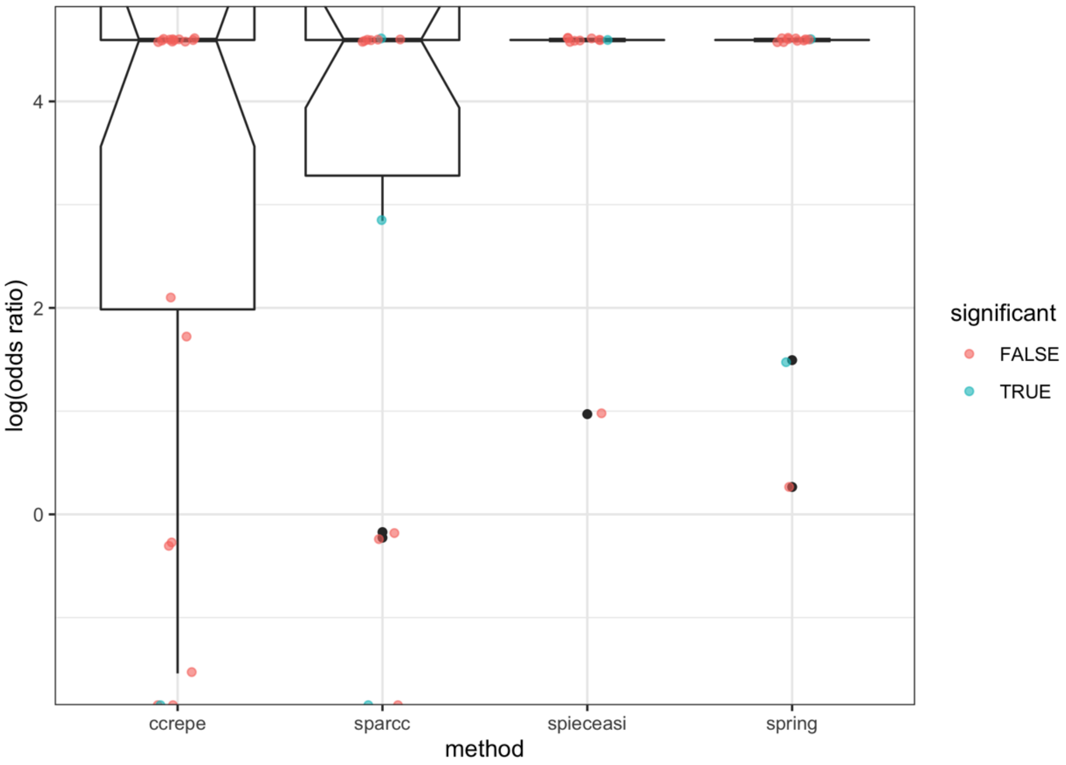


**Supplementary Figure 15**. Distribution of the logarithm (base10) of the odds ratio for the occurrence of a copresence association among genera belonging to the same family compared to genera belonging to different families. A dummy value of 5 was used when the odds ratio was infinite. The colour of the points shows if a Chi-square test of the null hypothesis that the frequency of copresence associations was not different between members of the same or different families (p<0.05).

**Supplementary Table 2.** The 50 most frequent associations detected at the species level using 4 inference methods (see section 2). For each edge n indicates the number of methods which detected the association and med_ebq the median quantile for edge betweenness. Ass. type indicates the nature of the association (copres = copresence, mut. ex. mutual exclusion). Some associations were detected in more than one data set

| dataset | ass.type | edge name | n | med_ebq | Reference |
|---|---|---|---|---|---|
| ST23 | copres | *Brevibacterium aureum--Staphylococcus equorum* | 4 | 0.94 | Bassi et al., 2015 |
| ST49 | copres | *Corynebacterium variabile--Levilactobacillus brevis* | 4 | 0.76 | Kamimura et al., 2019 |
| ST149 | copres | *Caryophanon latum--Porphyromonas levii* | 4 | 0.69 | Cremonesi et al., 2020 |
| ST49 | copres | *Psychrobacter sanguinis--Vibrio rumoiensis* | 4 | 0.68 | Kamimura et al., 2019 |
| ST22 | copres | *Staphylococcus equorum--Staphylococcus sciuri* | 4 | 0.60 | Calasso et al., 2016 |
| ST49 | copres | *Staphylococcus equorum--Staphylococcus sciuri* | 4 | 0.59 | Kamimura et al., 2019 |
| ST9 | copres | *Lactobacillus delbrueckii--Lactobacillus helveticus* | 4 | 0.58 | Alessandria et al., 2016 |
| ST41 | copres | *Marinomonas foliarum--Vibrio litoralis* | 4 | 0.52 | Dugat-Bony et al., 2016 |
| ST49 | copres | *Psychrobacter pacificensis--Psychrobacter sanguinis* | 4 | 0.45 | Kamimura et al., 2019 |
| ST49 | copres | *Marinilactibacillus psychrotolerans--Psychrobacter pacificensis* | 4 | 0.42 | Kamimura et al., 2019 |
| ST43 | copres | *Alkalibacterium gilvum--Marinilactibacillus psychrotolerans* | 4 | 0.38 | Guzzon et al., 2017 |
| ST49 | copres | *Ligilactobacillus acidipiscis--Weissella paramesenteroides* | 4 | 0.34 | Kamimura et al., 2019 |
| ST149 | copres | *Chryseobacterium anthropi--Chryseobacterium haifense* | 4 | 0.11 | Cremonesi et al., 2020 |
| ST41 | copres | *Vibrio litoralis--Vibrio rumoiensis* | 3 | 0.92 | Dugat-Bony et al., 2016 |
| ST48 | copres | *Brochothrix thermosphacta--Rahnella aquatilis* | 3 | 0.86 | Marino et al., 2019 |
| ST149 | copres | *Acinetobacter guillouiae--Chryseobacterium haifense* | 3 | 0.79 | Cremonesi et al., 2020 |
| ST48 | copres | *Anoxybacillus flavithermus--Shewanella putrefaciens* | 3 | 0.69 | Marino et al., 2019 |
| ST49 | copres | *Lactiplantibacillus paraplantarum--Levilactobacillus brevis* | 3 | 0.68 | Kamimura et al., 2019 |
| ST48 | copres | *Lactobacillus delbrueckii--Lactobacillus helveticus* | 3 | 0.67 | Marino et al., 2019 |
| ST18 | copres | *Propionibacterium acnes--Pseudomonas mandelii* | 3 | 0.66 | Stellato et al., 2015 |
| ST41 | copres | *Halomonas zhanjiangensis--Vibrio toranzoniae* | 3 | 0.65 | Dugat-Bony et al., 2016 |
| ST22 | copres | *Staphylococcus equorum--Streptococcus pneumoniae* | 3 | 0.65 | Calasso et al., 2016 |
| ST23 | copres | *Lacticaseibacillus rhamnosus--Lentilactobacillus buchneri* | 3 | 0.56 | Bassi et al., 2015 |
| ST3 | copres | *Lactococcus lactis--Streptococcus thermophilus* | 3 | 0.50 | De Filippis et al., 2014 |
| ST6 | copres | *Lactiplantibacillus pentosus--Latilactobacillus fuchuensis* | 3 | 0.50 | De Pasquale et al., 2014a |
| ST49 | copres | *Corynebacterium stationis--Psychrobacter meningitidis* | 3 | 0.47 | Kamimura et al., 2019 |
| ST49 | copres | *Corynebacterium stationis--Tetragenococcus halophilus* | 3 | 0.45 | Kamimura et al., 2019 |
| ST49 | copres | *Lactiplantibacillus paraplantarum--Weissella paramesenteroides* | 3 | 0.43 | Kamimura et al., 2019 |
| ST146 | copres | *Lactococcus raffinolactis--Streptococcus parauberis* | 3 | 0.38 | Castellanos-Rozo et al., 2020 |
| ST49 | mut_ex | *Lactiplantibacillus paraplantarum--Leuconostoc mesenteroides* | 3 | 0.34 | Kamimura et al., 2019 |
| ST49 | copres | *Tetragenococcus halophilus--Weissella paramesenteroides* | 3 | 0.32 | Kamimura et al., 2019 |
| ST22 | copres | *Corynebacterium tuberculostearicum--Propionibacterium granulosum* | 3 | 0.27 | Calasso et al., 2016 |
| ST49 | copres | *Acinetobacter johnsonii--Pseudomonas fragi* | 3 | 0.21 | Kamimura et al., 2019 |
| ST8 | copres | *Enterococcus faecalis--Macrococcus caseolyticus* | 3 | 0.21 | Dolci et al., 2014 |
| ST49 | copres | *Corynebacterium stationis--Corynebacterium variabile* | 3 | 0.18 | Kamimura et al., 2019 |
| ST49 | copres | *Ligilactobacillus acidipiscis--Tetragenococcus halophilus* | 3 | 0.16 | Kamimura et al., 2019 |
| ST9 | copres | *Lacticaseibacillus casei--Limosilactobacillus fermentum* | 3 | 0.05 | Alessandria et al., 2016 |

| | | | | | |
|---|---|---|---|---|---|
| ST10RNA | copres | *Lacticaseibacillus casei--Limosilactobacillus fermentum* | 2 | 1.00 | De Filippis et al., 2016 |
| ST8 | copres | *Lactobacillus delbrueckii--Streptococcus thermophilus* | 2 | 1.00 | Dolci et al., 2014 |
| ST1 | copres | *Lactococcus garvieae--Weissella viridescens* | 2 | 0.96 | Guidone et al., 2016 |
| ST18 | copres | *Lactobacillus delbrueckii--Lactobacillus helveticus* | 2 | 0.96 | Stellato et al., 2015 |
| ST18 | copres | *Chromohalobacter canadensis--Lactobacillus delbrueckii* | 2 | 0.88 | Stellato et al., 2015 |
| ST8 | mut_ex | *Acinetobacter johnsonii--Lactobacillus delbrueckii* | 2 | 0.84 | Dolci et al., 2014 |
| ST10RNA | copres | *Lactobacillus helveticus--Streptococcus thermophilus* | 2 | 0.83 | De Filippis et al., 2016 |
| ST23 | copres | *Enterococcus casseliflavus--Limosilactobacillus fermentum* | 2 | 0.82 | Bassi et al., 2015 |
| ST22 | copres | *Staphylococcus sciuri--Streptococcus pneumoniae* | 2 | 0.80 | Calasso et al., 2016 |
| ST10RNA | copres | *Lacticaseibacillus casei--Lentilactobacillus kefiri* | 2 | 0.79 | De Filippis et al., 2016 |
| ST22 | copres | *Clostridium difficile--Propionibacterium granulosum* | 2 | 0.78 | Calasso et al., 2016 |
| ST49 | copres | *Corynebacterium stationis--Psychrobacter celer* | 2 | 0.78 | Kamimura et al., 2019 |
| ST22 | copres | *Clostridium difficile--Lactiplantibacillus pentosus* | 2 | 0.77 | Calasso et al., 2016 |
